# Molecular co-accessibility identifies coordinated regulation between distant *cis*-regulatory elements

**DOI:** 10.1101/2025.08.05.668660

**Authors:** Mathias Boulanger, Kasit Chatsirisupachai, Rene Snajder, Marc Jan Bonder, Karine Lapouge, Kim Remans, Arnaud R Krebs

## Abstract

In metazoans, gene expression is typically regulated by a *cis*-regulatory landscape (CRL) composed of a promoter and multiple enhancers. How these *cis*-regulatory elements (CREs) coordinate their function across large genomic distances remains unclear. For example, is the simultaneous activation of multiple enhancers required to promote transcription? Here, we combined single-molecule footprinting with long-read sequencing to quantify how often chromatin accessibility and transcription factor binding co-occur across entire CRLs in the *Drosophila* genome. Analysis of thousands of individual DNA molecules at each locus revealed that CREs form a specific network with shared single-molecule chromatin accessibility profiles. Co-accessibility is not limited to adjacent CREs, and is frequently observed between CREs brought into proximity by chromatin looping. Co-accessible CREs exhibit strong coordination in their cell type-specific accessibility, linking enhancer activity with transcription activation. Our data uncover dependencies between CREs genome-wide, and suggest that coordinated enhancer activation is a widespread mechanism regulating gene expression.

## Introduction

*Cis*-regulatory elements (CREs) are short non-coding DNA stretches that contain clusters of binding sites for transcription factors (TFs). Binding of TFs at CREs leads to the opening of chromatin and the subsequent recruitment of RNA Polymerase II on DNA to initiate transcription. Each gene is regulated by a promoter and several enhancers that form its *cis*-regulatory landscape (CRL). Enhancers are located kilobases to megabases away from the gene they regulate (1, 2). They act through formation of chromatin loops to come in physical proximity of the promoter and the gene they activate (3, 4).

Activity of many genes, including most developmental genes, is regulated by multiple enhancers. Several models have been proposed to explain how they combine their function (reviewed in Uyehara and Apostolou, 2023 (2)). The simplest model suggests that enhancers act as individual units, with redundant or additive function on gene expression (5–9). For instance, enhancer deletion studies have supported the additive model where multiple enhancers are required to reach sufficient levels of expression of the target gene, or to precisely modulate its expression at different times of development (5, 6, 8, 9). Several pieces of evidence argue for a more complex model where the combined function of multiple enhancers goes beyond the simple sum of their individual function. For instance, identification of regulatory loci with unusually high levels of cofactor recruitment suggested the existence of higher order cooperativity between clusters of enhancers (10, 11). This is further supported by the observation of non-additive effects in enhancer deletion studies (12, 13). Moreover, several studies reported the existence of distal CREs that have weak intrinsic transcription activity, but are indispensable to potentiate the activity of other enhancers at the locus (14–16), indicating a functional specialisation of CREs within CRLs. Together, this suggests the existence of a complex network of dependencies between enhancers and the potential requirement for their coordinated function to activate transcription.

Coordination in the activity of CREs has been primarily observed at the level of promoters. For instance, live imaging of transcription of two fluorescent reporters showed that the same enhancer can activate two promoters simultaneously (17). Transcriptional coupling has been further observed for paralogous genes in *Drosophila* that are frequently looping into the same 3D environment, and show coordinated transcription (18–20). Evidence of coordinated transcription has also been suggested to take place for functionally related genes in mammalian cells (21). Whether the multiple enhancers and the promoter involved in the activation of the same gene are coordinated in their activation remains an important open question.

Mapping TF binding, chromatin accessibility or the presence of chromatin marks is frequently used to identify active enhancers (22). This is achieved through bulk genomics assays that measure the average regulatory activity at individual CREs across large cell populations, thus not informing on the potential coordination in their activation (23, 24). Spatial proximity between CREs can be used to study functional interactions between CREs (3, 4). Yet, a large fraction of 3D loops between CREs are disconnected from regulatory activity (25–28), challenging the use of spatial proximity to unambiguously establish coordinated regulation within a CRL. Single-molecule genomics technologies have recently emerged as a strategy to study dependencies between regulatory events in the genome (23, 24, 29). These assays are based on methylation footprinting of chromatin accessibility, that preserves the integrity of DNA, allowing quantification of multiple regulatory events on the same molecules (23, 24). The frequency at which two regulatory events co-occur on the same DNA molecules can subsequently be used to define positive and negative dependencies between them (23). For instance, we used single-molecule footprinting (SMF) to show that molecular co-occupancy identifies cooperativity between TFs (30). Conversely, we found that molecular exclusion can identify repression of enhancer activity by DNA methylation (31).

Coupling methylation footprinting with long-read sequencing technologies measures chromatin accessibility continuously over molecules of up to hundred kilobases, allowing to detect the molecular co-occurrence of regulatory events across long genomic distances (32–36). A prerequisite for the quantification and statistical interpretation of molecular co-occurrence is to sequence hundreds of molecules that cover the two loci of interest (37). This has been achieved for the compact genome of *Saccharomyces cerevisiae* (35, 38), but where enhancers are absent and gene expression is primarily controlled at the level of promoters. In metazoans, genes are controlled by multiple enhancers and genomes are larger by several orders of magnitude, challenging the systematic identification of co-occurring regulatory events across CREs in existing datasets (20, 34, 36, 39).

In this study, we took advantage of the compact *Drosophila* genome, where most enhancers lie within 6 kb of their target promoters (26, 40), to systematically measure if chromatin opening is coordinated across multiple enhancers of a CRL. We coupled SMF with long-molecule sequencing in two cell lines with divergent transcriptional programs. We sequenced these samples at a depth compatible with the statistical quantification of coordinated chromatin opening across entire CRLs genome-wide. The resulting maps suggest that at each locus, a specific network of enhancers and promoters are co-accessible. Co-accessibility typically involves multiple CREs within a CRL, extending beyond adjacent CREs and involving CREs loci distant by up to 29 kb. These CREs are frequently brought in spatial proximity through 3D chromatin looping, and have molecular co-occupancy of insulator proteins. Coordinated changes in cis-regulatory activity across co-accessible enhancers confirm their functional association and links them to the transcriptional response at their target gene.

## Results

### Quantifying simultaneous chromatin accessibility of entire *cis*-regulatory landscapes

We set out to measure the frequency at which chromatin accessibility co-occurs between the multiple enhancers and promoters that form a CRL. We adapted principles of single-molecule long-read accessible chromatin mapping sequencing (SMAC-seq) (35), where accessible DNA is footprinted using recombinant cytosine methyltransferases and methylation is directly called from reads of several kilobases that are generated by Oxford Nanopore Technology (ONT) (Figure 1A). Similar to SMF (37, 41), we used cytosine methyl-transferases (GpC – M.CviPI and CpG – M.SssI), for which methylation calls using ONT are more reliable than those in adenine contexts (35). We trained a custom methylation caller for these two dinucleotide contexts, improving existing models trained only on CpGs (Figure S1A-D). This resulted in methylation profiling accuracy comparable to reference SMF data obtained using bisulfite sequencing (41) (R > 0.85, Figure S1BC, E). The resulting SMF-ONT data captures chromatin accessibility, nucleosome, and TF footprints at single cytosine resolution on individual DNA molecules (exemplified in Figure 1B-C, Figure S1E).

**Fig. 1.**
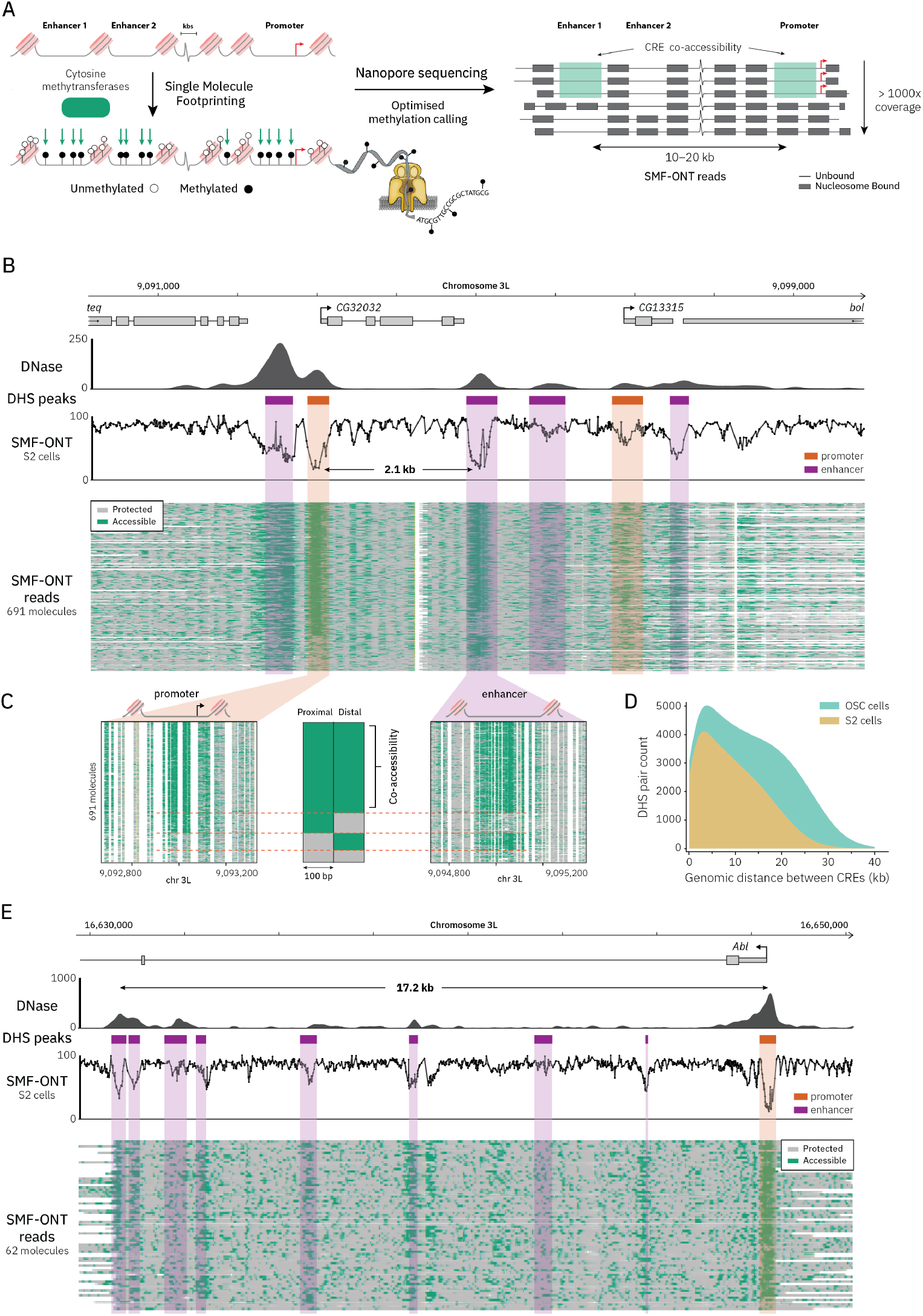
Simultaneous detection of chromatin accessibility across entire *cis*-regulatory landscapes. **(A)** Schematic overview of the approach based on single-molecule footprinting (SMF) coupled to direct sequencing using Oxford Nanopore Technology (ONT). Nuclei are extracted and DNA is footprinted using exogenous methyltransferases which methylate accessible cytosines in GpC and CpG dinucleotide contexts. High molecular weight DNA is extracted and sequenced using ONT, providing continuous chromatin accessibility measure across 10–20 kilobases. Applying this method to the compact genome of *Drosophila* generates datasets with a coverage compatible with the statistical analysis of the co-occurrence of chromatin accessibility across the majority of enhancer-promoter pairs. **(B)** Single-locus example illustrating simultaneous detection of chromatin accessibility at 6 CREs over 691 molecules. Genomic tracks are shown from top to bottom: DNase-seq signal, DHS peaks (orange: promoter; purple: putative enhancer), and average SMF signal (1 *−* methylation (%)) of individual cytosines. The bottom track displays individual SMF-ONT reads aligned to the region. Chromatin accessibility is colour-coded (grey: unmethylated—protected; green: methylated—accessible). **(C)** Zoomed-in view illustrating the co-accessibility between the promoter of the *CG32032* gene and a putative enhancer located 2.1 kb downstream. Left and right panels display the same individual SMF reads over the two regions, same colour code as in **(B)**. The middle panel summarises the accessibility status of each CRE (grey: protected from methylation; green: accessible). **(D)** Density plot showing the distribution of the distances between CREs covered in our datasets in S2 (yellow) or OSC cells (green). **(E)** Single-locus example illustrating the simultaneous measure of chromatin accessibility at a promoter and 8 putative enhancers scattered over 20 kb. Same representation as **(B)**.

Quantifying co-occurrence of two regulatory events requires sampling hundreds of molecules that span both loci. Here, we chose *Drosophila melanogaster* as a model where most genes are regulated by a complex CRL composed of a promoter and multiple enhancers; ii) the median distance between promoters and its closest enhancer is 6 kb, facilitating their coverage within the same sequencing read; iii) the genome size is an order of magnitude smaller than that of mammals, allowing the sequencing of thousands of DNA molecules of each individual locus (Figure S1F).

We generated high-coverage data (> 1,000x) in two cell lines with embryonic and somatic gene expression programs (S2 and Ovarian Somatic Cells (OSC)) (Figure S1F). The resulting dataset is an order of magnitude larger than existing single-molecule chromatin accessibility datasets (160x Fiberseq datasets in *Drosophila* Schneider (S2) cells (20, 36)). The data reproducibly capture chromatin accessibility in each cell type, as well as differential chromatin accessibility at cell type-specific CREs (Figure S1G-I) (41, 42).

In order to quantify co-accessibility between CREs, we first annotated active CREs in our system based on chromatin accessibility using existing DNase I hypersensitive sites (DHSs) (42). We categorised these according to their proximity to TSSs, separating promoters from putative enhancers (see Methods). Proximal and distal accessible sites showed expected enrichment for specific chromatin mark combinations for promoters and enhancers. Additionally, distal accessible sites were enriched for enhancer activity in parallelised reporter assays (STARR-seq, Figure S2A-D) (42).

The coverage of our datasets is sufficient to collect several hundreds of simultaneous measurements of chromatin accessibility across multiple CREs on individual DNA molecules. For instance, we obtained 691 molecules covering the promoter region of the *CG32032* gene and a putative enhancer located 2 kb downstream (Figure 1B, bottom panel). We could distinguish DNA molecules where both CREs were simultaneously accessible, from those where chromatin accessibility is detected at either of them (Figure 1C). The number of molecules spanning distant CREs drops as a function of the genomic distance between them (Figure S1F). With an average fragment size of 11 kb, we could resolve chromatin accessibility simultaneously for 68,000 CRE pairs separated by a median distance of 8 kb in S2 cells (Figure 1D, Figure S1F). This is sufficient to resolve the accessibility status of entire CRLs, such as the one of the *Abl* gene (Figure 1E). This CRL spans 17 kb and contains a promoter and 8 putative enhancers that are all simultaneously resolved within individual molecules (Figure 1E, Figure S1J). In summary, we generated genome-wide single-molecule chromatin accessibility datasets with sufficient depth to systematically quantify the co-accessibility of promoters and multiple enhancers on the same DNA molecules.

### Co-accessibility occurs between specific sets of CREs

We next quantified the frequency at which enhancers and promoters are accessible on individual DNA molecules, across the genome (Figure S2E, see Methods). We observed that most active CREs are accessible only in a fraction of the sampled molecules (Figure 2A). This heterogeneity of chromatin accessibility is particularly pronounced for putative enhancers, in agreement with previous reports (36, 43). We focused on CREs that show heterogeneous accessibility in our system and measured the frequency at which they are co-accessible on the same DNA molecules. We performed a Fisher’s exact test to assess the independence in chromatin opening between each pair of CRE sufficiently covered in our assay (Figure 2B, see Methods). Based on this test, we separated co-accessible pairs of CREs, from independent pairs or pairs that show exclusive accessibility (Figure 2C, Figure S2F, see Methods). We were able to test co-accessibility of each CRE against 8 other CREs in median, with up to 29 neighbouring CREs at certain loci (Figure S2G). We ensured that the effect size of the test (odds ratio) and its statistical significance (*p*-value) were not skewed by the level of accessibility of the CREs being tested (Figure S2H-J). We identified 12,901 pairs of CREs which have more co-accessibility than expected by chance in S2 cells. This contrasts with the majority of the tested pairs (54,774), that show coaccessibility levels compatible with their random occurrence (Figure 2C, Figure S2F). Similar proportions of co-accessible CREs were detected in OSC cells (Figure S2F). This suggests that the opening of chromatin can be coordinated in cis, and that this is restricted to a subset of CREs at each locus. This specificity is evident when visualizing the results of the coaccessibility test across the genome (Figure 2D). While 78% of the tested CREs showed co-accessibility with at least one neighbouring element, most remained independent from the majority of their neighbours (Figure 2D). The differences between co-accessible and independent CREs can be observed when analysing single-molecule accessibility patterns for individual pairs (Figure 2E-G). For example, the *Hsp22* promoter (promoter 2, Figure 2E) is co-accessible with 4 CREs including the *Fdx1* promoter, where a large fraction of the molecules has co-accessibility (Figure 2E-F). In contrast, the same promoter has simultaneous chromatin accessibility on a smaller fraction of molecules with an enhancer located 6.5 kb downstream (Pair 2, Figure 2E, 2G). This suggests that coaccessibility occurs between specific sets of CREs at each locus, potentially identifying functional relations within a CRL.

**Fig. 2.**
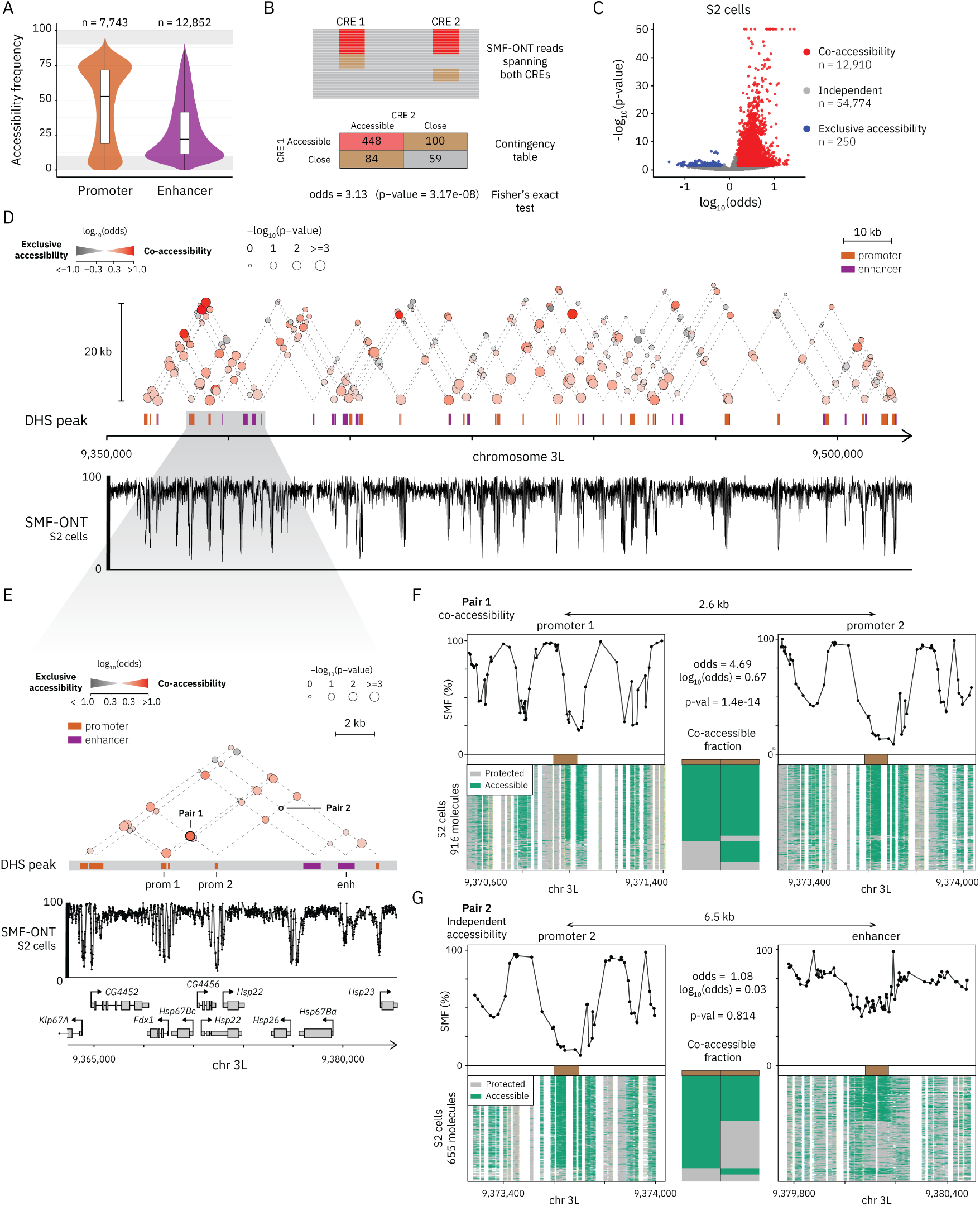
Quantification of the molecular co-accessibility across distant CREs. **(A)** Violin box plots comparing the distribution of the frequencies of chromatin accessibility at promoters (orange) and putative enhancers (purple). The middle line of the box represents the median. The box displays the interquartile range (IQR), 25^th^ to 75^th^ percentile. Whiskers represent a distance of 1.5 × IQR. Violin shapes depict the density of data points at each value. The number of CREs in each category is indicated above the plot. **(B)** Schematic representation of the strategy used to quantify co-accessibility across CREs. CREs were defined using peaks of DNase I hypersensitive sites (DHSs). Sequencing reads were sorted according to the chromatin accessibility measured within a 100 bp window centred around the peak. For each CRE pair, individual molecules were categorised as co-accessible on both CREs (red), accessible at only one CRE (brown), or inaccessible at both CREs (grey). The interdependency of chromatin accessibility between CREs was tested using Fisher’s exact test over the resulting contingency table. We used a *p*-value < 0.05 and log_10_(odds ratio) *>* log_10_(1.5) as a threshold for statistical significance of co-accessibility. **(C)** Volcano plot illustrating the results of co-accessibility tests in S2 cells. Colours represent the result of the test (red: co-accessibility; blue: exclusive accessibility; grey: independent). **(D)** Pairwise map of co-accessibility across a 150 kb example region. Each dot represents the co-accessibility between two pairs of CREs that are joined by dotted lines. The colour of the dots represents the odds ratio from the Fisher’s exact test (grey: exclusive accessibility; red: co-accessibility). The size of the dot represents the *p*-value of the test. The lower genomic tracks show average SMF signal (1 *−* methylation (%)) of individual cytosines and DHS peaks (orange: promoter; purple: putative enhancer). **(E)** Zoomed-in view illustrating a pair-wise map of co-accessibility at 15 kb region. Same representation as **(D). (F)** Example of a co-accessible CRE pair, separated by 2.6 kb (Pair 1, Promoter 1 and Promoter 2 in **(E)**). The upper panel shows the average SMF signal in a 1 kb window around each CRE. The brown box marks the bin used to collect cytosine methylation data. The lower panel shows single-molecule stacks across both CREs (916 molecules). Each row corresponds to the same DNA molecule for both CREs. Protected cytosines are displayed in grey, while accessible cytosines are shown in green. The middle bars depict the frequency of each state (grey: protected; green: accessible). Odds ratio and *p*-value from the Fisher’s exact test for co-accessibility are shown. **(G)** Example of a CRE pair with independent accessibility, separated by 6.5 kb (Pair 2, Promoter 2 and Enhancer 1 in **(E)**). Same representation as **(F)**.

### Co-accessibility is not restricted to adjacent CREs

We next analysed the nature and the genomic distribution of the CREs where co-accessibility is detected. We first asked if co-accessibility preferentially occurs at enhancer-promoter pairs that are expected to communicate to regulate gene expression. We indeed observed co-accessibility for 34% of the enhancer-promoter pairs tested in S2 cells (Figure 3A), and similar proportions in OSC cells (Figure S3A). Yet, coaccessibility is not restricted to this type or CRE, as similar proportions of enhancer-enhancer pairs, and an even larger fraction of promoter-promoter pairs showed co-accessibility (Figure 3A, Figure S3A).

**Fig. 3.**
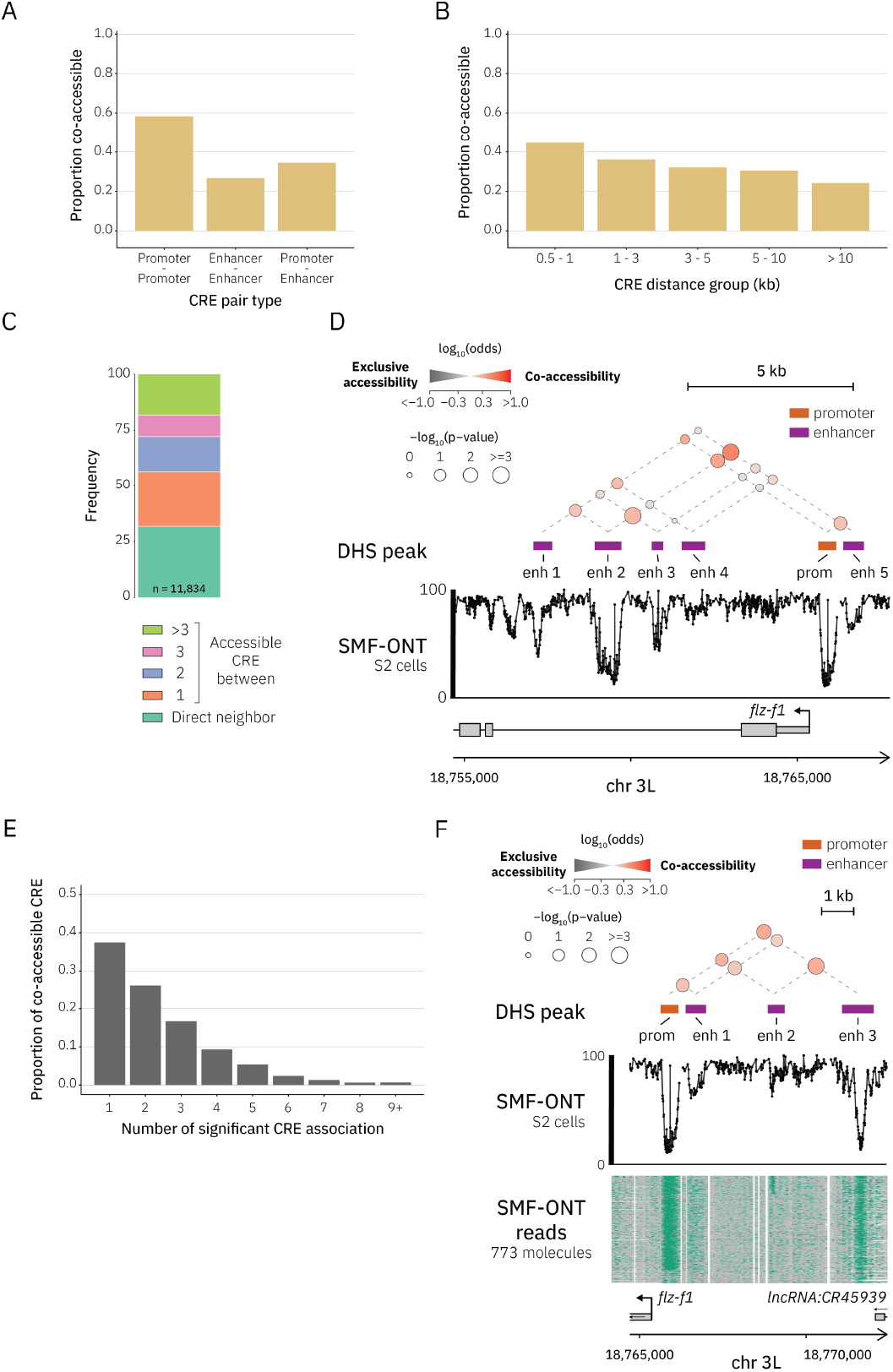
Genomic distribution of CREs with molecular co-accessibility. **(A)** Co-accessibility is not restricted to enhancer-promoter pairs. Bar chart displaying the proportion of co-accessible CRE pairs categorised by pair type in S2 cells. **(B)** Co-accessibility occurs at distant CREs. Bar charts illustrating the proportion of co-accessible CRE pairs as a function of genomic distance between them. Only pairs with > 500x coverage in S2 cells were included in this analysis to avoid false negatives due to low coverage. **(C)** Co-accessibility is not restricted to adjacent CREs. Stacked bar plots depicting the number of CREs intercalated between two co-accessible CREs in S2 cells. **(D)** Pair-wise map of co-accessibility illustrating the distribution of co-accessible CREs over the *flz-f1* locus. Same representation as Figure 2D. **(E)** Individual CREs can be co-accessible with multiple neighbours. Histogram displaying the number of significant co-accessible pairs for each CRE. **(F)** Pair-wise map of co-accessibility at a cluster of co-accessible CREs. Single-locus example showing the promoter region of the *flz-f1* gene, which is co-accessible with three putative enhancers. Same representation as Figure 2D and 2F-G.

We next wondered if CREs that are located close to one another are more likely to be co-accessible. We plotted the proportion of co-accessible CRE pairs as a function of the distance between them (Figure 3B). To avoid confounding effects linked to the drop in sequencing depth at larger genomic distances, we only considered pairs with saturating coverage in this analysis (> 500x, Figure S2K, see Methods). We observed that the proportion of pairs showing coaccessibility decreased as a function of distance (Figure 3B, Figure S3B). Yet, > 24% of the pairs separated by 10 kb were co-accessible, and we could detect co-accessibility of CREs separated by over 24 kb. We note here that the size distribution of our sequencing reads limits our analysis to < 40 kb, and that co-accessibility that may occur at larger genomic distances is not detectable with this data. Only 32% of the co-accessible pairs occur between CREs adjacent to one another, with the majority of co-accessible events occurring between those separated by one or more intervening CREs (Figure 3C, Figure S3C-D). This is, for instance, the case for the *flz-f1* promoter (Figure 3D), that has independent accessibility with two adjacent enhancers (Figure 3D, enhancer 3-4), but shows significant co-accessibility with one enhancer located at larger genomic distance (Figure 3D, enhancer 1-2).

A promoter is typically regulated by multiple enhancers. To investigate whether co-accessibility extends beyond pairs of CREs, we asked whether individual CREs can be coaccessible with multiple others, and whether such interactions are spatially clustered in the genome. Over 60% of co-accessible CREs were linked to more than one partner, with some connected to as many as 14 others (Figure 3E, Figure S3E). While co-accessible CREs are often dispersed across the locus (Figure 2D, Figure 3D), we also identified regions where several neighbouring CREs were co-accessible (Figure 3F). These findings suggest that specific sets of enhancers and promoters are frequently accessible simultaneously on the same DNA molecules.

### Su(Hw) and GAGA-factor bind co-accessible CREs

Since co-accessibility is restricted to a specific set of CREs, we hypothesised that certain TFs may be directly or indirectly involved in regulating their activity. We tested if the binding motifs of certain TFs may be enriched at co-accessible CRE pairs over independent pairs. Unlike classical onesided motif enrichment analysis, we tested here for the simultaneous presence of motifs on both CREs of a pair (see Methods, Figure 4A). We identified 248 out of 11,026 motif pairs to be specifically enriched at co-accessible CRE pairs in S2 cells (Figure 4B-C, Figure S4A). These enriched motifs entail those bound by ubiquitously expressed insulator proteins, such as BEAF-32 and Su(Hw), as well as developmental factors such as Pnr and Ara (Figure 4B-C, Figure S4A). We also found an enrichment for Trithorax-like (Trl), also known as GAGA factor (GAF), that has recently been found to be important for enhancer-promoter communication (Figure 4B-C, Figure S4A) (44). We next asked if these motifs were enriched in specific types of CRE pairs (Figure 4C, Figure S4A). We found that Su(Hw) and Trl motifs were primarily found at enhancer-enhancer and enhancer-promoter pairs, while other motifs were almost exclusively found at promoter-promoter pairs (Figure 4C, Figure S4B-D). These enrichments are not cell-type specific as similar sets of motifs were found enriched in OSC (Figure S4E-G).

Leveraging the ability of SMF to detect TF footprints (Figure S1E) (30), we next asked if we can observe simultaneous binding of TFs on the same DNA molecules and if this may explain co-accessibility of certain pairs of CREs. We measured TF footprints on individual molecules genomewide, calculated their frequency of co-occurrence and tested the dependency between them (see Methods). About half of the Su(Hw)-Su(Hw) pairs located among co-accessible CREs showed significant co-occupancy (Figure 4D). In contrast, the majority of the binding events for other tested TF pairs located within co-accessible CRE pairs were independent (Figure 4D). This high Su(Hw) co-occupancy across distant CREs can be observed at individual loci as exemplified in Figure 4E. We noted however that the frequency of co-binding across distant loci was much lower than when the two motifs are contained within the same regulatory elements (Figure 4F). This was further evident when contrasting Su(Hw) co-occupancy within a CRE where 84% of the molecules are co-bound (Figure 4G) with binding across CREs where only 30% of the molecules are co-bound (Figure 4E). Together this suggests that specific TFs may be involved in regulating co-accessibility, and that some of them show molecular co-occupancy across distant loci.

**Fig. 4.**
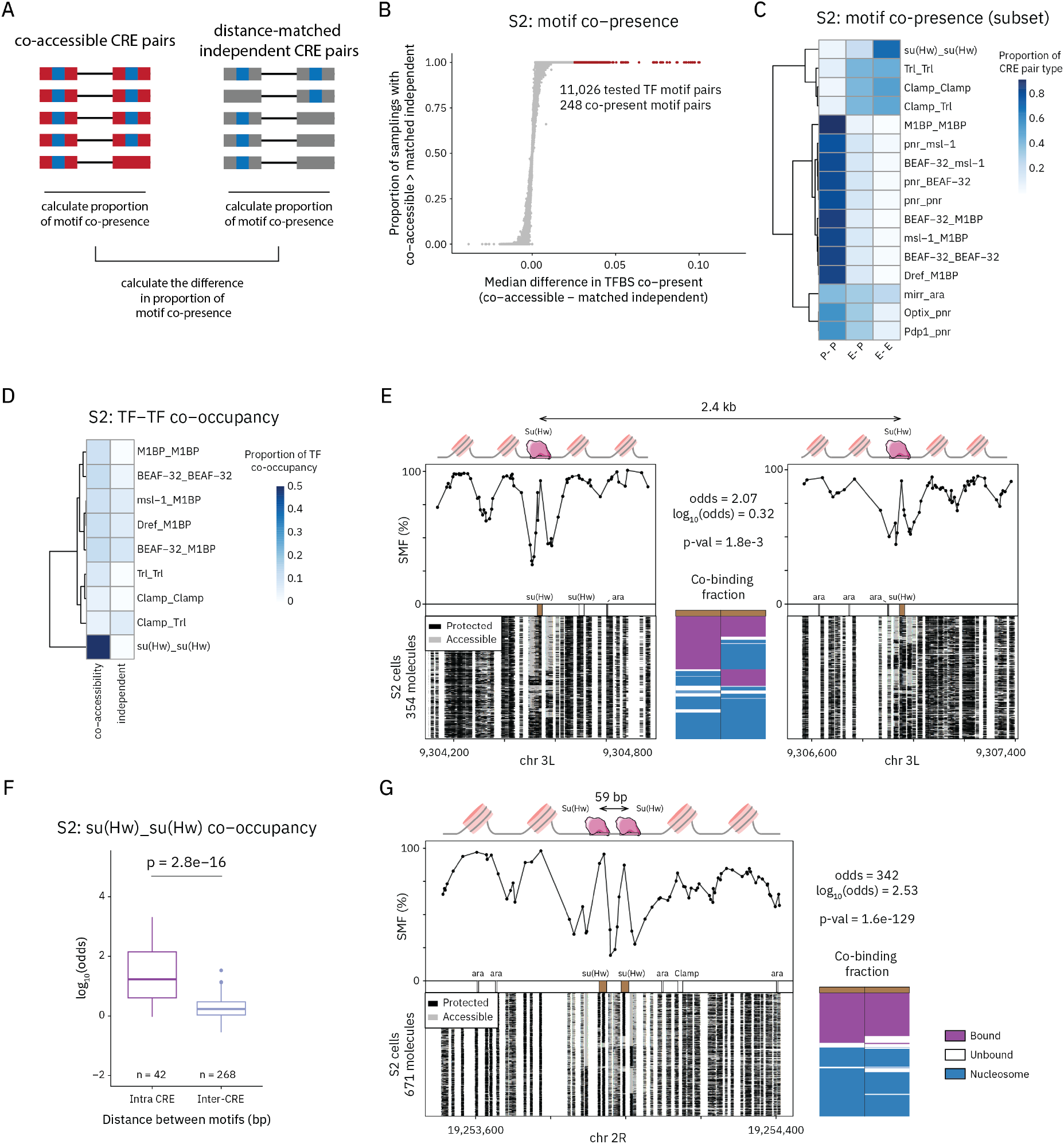
Transcription factor binding at co-accessible CRE pairs. **(A)** Schematic representation of the method used to test for simultaneous presence of transcription factor (TF) binding motifs in co-accessible CREs. For each pair of motifs, the proportion of CREs with both motifs present was calculated in co-accessible and independent CRE pairs (see Methods). **(B)** Specific TF motifs are enriched at co-accessible CREs. Scatter plot displaying the enrichment for each TF motif pair in co-accessible versus independent CRE pairs in S2 cells. The x-axis represents the median difference in the proportion of pairs with motif co-presence between co-accessible and independent pairs. The y-axis shows the proportion of samplings in which co-accessible pairs exhibit a higher motif co-presence than distance-matched independent pairs. Top candidate motif pairs with the highest median difference are highlighted in red. **(C)** Different motifs are enriched at promoters and enhancers. Heatmap showing the distribution of the top motif pairs enriched as a function of the type of CREs involved. The colour scale represents the proportion of each CRE pair type (promoter-promoter, enhancer-promoter, enhancer-enhancer) in which the motif pairs are located. For the full heatmap, see Figure S4A. **(D)** Molecular co-occupancy of TFs is observed at co-accessible CREs. Heatmap comparing the proportion of CREs with significant TF-TF co-occupancy for co-accessible and independent pairs. **(E)** Single locus example of TF co-occupancy for Su(Hw) at CREs separated by 2.4 kb. Upper panel shows the average SMF signal in a 1 kb window around the Su(Hw) binding motifs (brown box). Lower panel shows single-molecule stacks across both regions (354 molecules, grey: accessible; black: inaccessible). The same DNA molecules are displayed on both panels. The middle barplot depicts the frequency of each state (purple: bound; white: unbound; blue: nucleosome). Odds ratio and *p*-value from the Fisher’s exact test for TF co-occupancy are shown. **(F)** Molecular co-occupancy between distant CREs is low compared to co-occupancy within CREs. Boxplot comparing TF co-occupancy within (< 200 bp: intra-CRE) and across CREs (≥ 200 bp: inter-CRE) as measured by the log_10_(odds ratio) from Fisher’s exact tests. The middle line of each box represents the median; boxes indicate the interquartile range (IQR; 25^th^ to 75^th^ percentile); whiskers extend to 1.5 × IQR. Statistical comparisons between groups were performed using the Wilcoxon rank-sum test. **(G)** Single locus example of high TF co-occupancy for Su(Hw) within the same CRE. Same representation as **(E)**.

### Co-accessible CREs have higher chromatin contact frequency

Since co-accessibility is not explained by genomic proximity, we wondered if 3D genome interactions may contribute to bringing co-accessible CREs in physical proximity. We therefore tested whether co-accessible CRE pairs are engaged in looping interactions using existing Micro-C data (45). We focused our analysis on CREs distant by 5–10 kb. This distance is enough to reduce biases due to proximity ligation in Micro-C, but close-enough to measure co-accessibility of many CREs with sufficient coverage. We compared 3D contact frequencies between independent and co-accessible pairs and found that co-accessible pairs showed more frequent 3D contacts than the independent ones (Figure 5A). This enrichment for 3D contacts at co-accessible CREs was consistent across various genomic distances (Figure S5A). Similar conclusions can be drawn when overlaying 3D contacts with co-accessibility at individual loci (Figure 5B). For instance, the promoter of the shn gene exhibited co-accessibility and a high Micro-C contact with an enhancer located 12 kb downstream (Pair 1, Figure 5B). Despite being located closer, it showed independent chromatin accessibility and lower contact frequency with an intercalated enhancer (Pair 2, Figure 5B). Together, this suggests that coaccessibility occurs at CREs that are frequently in physical proximity in the nucleus.

**Fig. 5.**
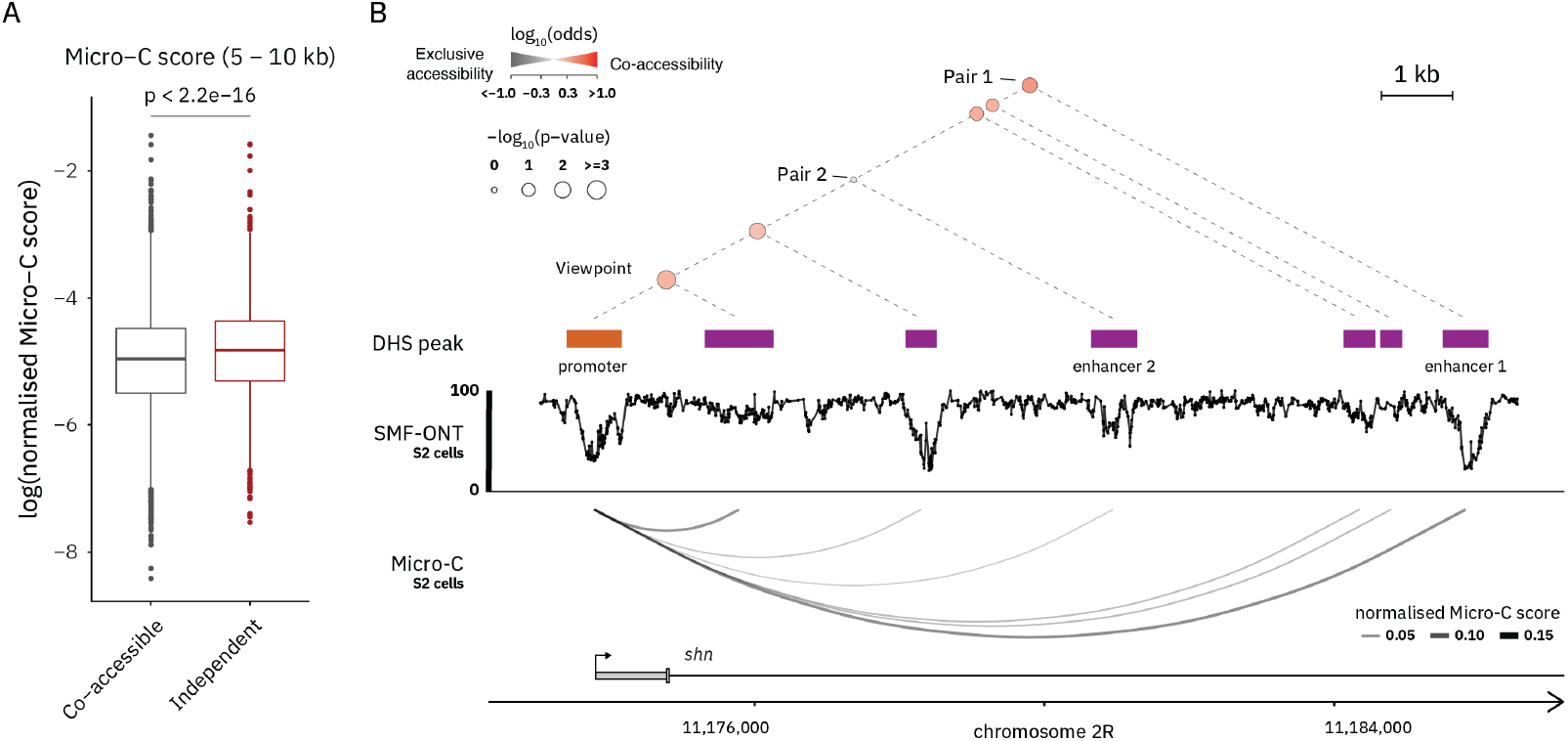
Relation between molecular co-accessibility and 3D chromatin architecture. **(A)** Co-accessible pairs have higher contact frequency than independent pairs. Boxplots representing the distribution of normalised Micro-C contact frequency scores between CRE pairs, comparing independent (grey) and co-accessible (red) 5–10 kb pairs. The middle line of the box represents the median. The box displays the interquartile range (IQR), 25^th^ to 75^th^ percentile. Whiskers represent a distance of 1.5 × IQR. Statistical comparisons between groups were performed using the Wilcoxon rank-sum test. **(B)** Pairwise map of co-accessibility illustrating the relation between co-accessibility and chromatin contacts measured by Micro-C. The upper panel is presented in the same representation as Figure 2D. The lower panel shows normalised Micro-C contact scores, represented by the size and colour intensity of connecting lines. The co-accessible pair between the promoter and enhancer 1 (Pair 1), separated by 12 kb, shows a higher Micro-C score than the independent pair between the promoter and enhancer 2 (Pair 2), which are 7 kb apart.

### Co-accessibility identifies functional dependencies between CREs

The co-accessibility observed across distant CREs suggests that their regulatory activity may be coordinated. If correct, their chromatin accessibility should also co-vary upon changes in the *cis*-regulatory activity across cell types. Conversely, activity of independent CREs should not be coordinated (Figure 6A). To test this, we analysed differences in chromatin accessibility at enhancers, the most variable class of CREs, across cell types. We ensured that differences in the distance distribution between co-accessible and independent pairs would not bias our results, by sampling independent pairs to match the distance distribution of the co-accessible pairs (Figure S6A-B). We found that changes in cell-type specific chromatin accessibility were not correlated at independent enhancer-enhancer pairs (Figure 6B, Figure S6C). In contrast, co-accessible enhancer-enhancer pairs showed correlated changes in accessibility (Figure 6C, Figure S6C). Consistently, concordance in chromatin accessibility changes is significantly enriched at co-accessible enhancer pairs compared to independent pairs (Figure 6D, Figure S6D). We next investigated if the changes in chromatin accessibility at enhancers led to concordant changes at promoters that were linked based on co-accessibility. A large fraction of co-accessible enhancer-promoter pairs had correlated changes in their chromatin accessibility across cell types (Figure 6E-G, Figure S6E-F). This observation extends to co-accessible promoter-promoter pairs, yet with lower correlation (Figure S6G-K). The contrast in the co-variation of accessibility between independent and co-accessible CREs can be observed within individual CRLs (Figure 6H). For instance, we identified a putative cell type-specific enhancer with independent chromatin accessibility to the adjacent *roX1* promoter (Pair 1, enhancer-promoter 1, Figure 6H), but co-accessible with the promoter of *CG2930* gene (Pair 2, enhancer-promoter 2, Figure 6H). Upon opening of chromatin at this enhancer, we observed concomitant increase in accessibility at the *CG2930*, but not at the *roX1* promoter (Figure 6H). Together, our results suggest that coaccessibility identifies CREs that are coordinated in their regulation.

**Fig. 6.**
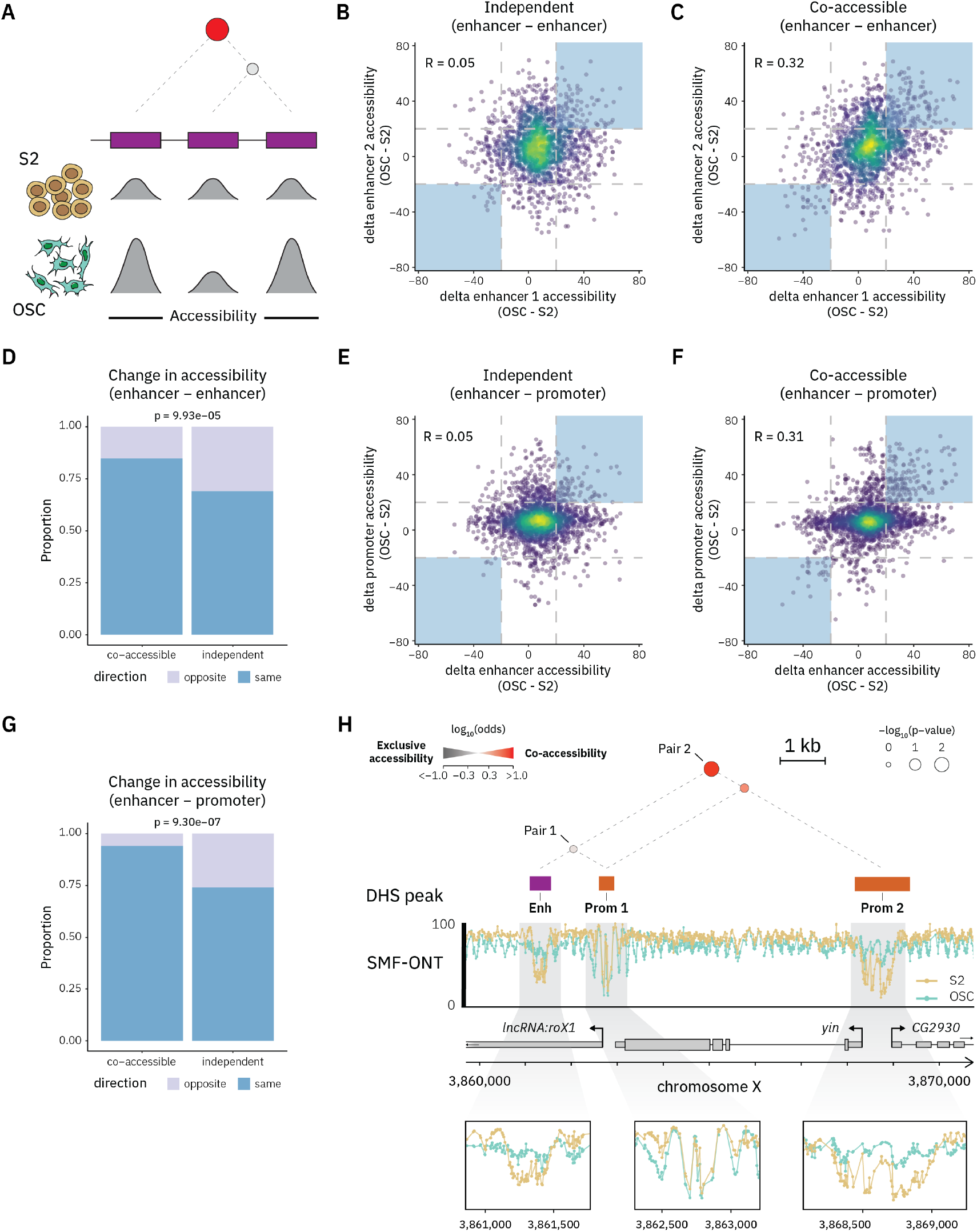
Coordinated cell type-specific changes in chromatin accessibility at co-accessible CREs. **(A)** Schematic representation of the hypothesis tested. If coaccessibility identifies dependencies between CREs, then chromatin accessibility should co-vary between cell types for co-accessible CREs, but not independent ones. **(B-C)** Cell type-specific chromatin accessibility changes are coordinated at co-accessible pairs of enhancers. Scatter plot showing the pairwise difference in chromatin accessibility between cell types at **(B)** independent and **(C)** co-accessible enhancer-enhancer pairs based on SMF. Independent pairs were sampled to be distance-matched to the co-accessible pairs. Blue shaded boxes highlight enhancer-enhancer pairs exhibiting highly coordinated changes in accessibility (> 20%). Pearson correlation coefficient (R) is displayed in the plot. **(D)** Stacked bar plot comparing the proportion of enhancer-enhancer pairs with coordinated chromatin accessibility changes across cell types. Only pairs with high variation in the accessibility of both enhancers (> 20%) were included. Statistical comparisons between groups were performed using a one-sided Fisher’s exact test. **(E-F)** Cell type-specific chromatin accessibility changes are coordinated at co-accessible enhancer-promoter pairs. Scatter plot showing the pairwise difference in chromatin accessibility between cell types at **(E)** independent and **(F)** co-accessible enhancer-promoter pairs. Same representation as **(B). (G)** Stacked bar plot comparing the proportion of enhancer-promoter pairs with coordinated chromatin accessibility changes across cell types. Same representation as **(D). (H)** Pair-wise map of co-accessibility illustrating co-variation of chromatin accessibility at co-accessible enhancer-promoter pairs. Pair 1 (enhancer and promoter 1) was found to be independent and does not exhibit coordinated changes in chromatin accessibility. In contrast, the chromatin accessibility of the co-accessible pair 2 (enhancer and promoter 2) shows coordinated increase between OSC and S2 cells. Same representation as Figure 2D.

### Molecular co-accessibility links enhancers activity to changes in gene expression

We next asked whether molecular co-accessibility could identify the functional dependencies within the CRLs that drive differential gene expression between cell types. We examined if changes in chromatin accessibility, or activity of distal enhancers would predict changes in the transcription of the target gene (Figure 7A). At co-accessible enhancer-promoter pairs, gain or loss of chromatin accessibility at the enhancer were significantly enriched for up or down-regulation of the target gene, respectively (Figure 7B, Supplementary Figure S7A). This effect is also observable when measuring changes in enhancer activity by STARR-seq (42). Changes in enhancer activity at co-accessible enhancer-promoter pairs better predicts the directionality of the changes in gene expression (Figure 7C-D, Figure S7B), compared to independent pairs. These functional associations could also be observed at the level of individual enhancers. For example, in OSCs, the enhancer located within the intron of the *alphas-Est9* gene is co-accessible with the *alphas-Est9* promoter located 6.5 kb downstream (Pair 1, enhancer-promoter 1, Figure 7E), but independent with the *alphas-Est10* promoter located 6.9 kb upstream (Pair 2, enhancer-promoter 2, Figure 7E). This enhancer is more accessible and active in OSC. Consistently, transcription of the *alpha-Est9* gene, but not the *alpha-Est10*, is up-regulated in OSCs (Figure 7E). Together, these results suggest that molecular co-accessibility accurately associates differential activity of enhancers with their functional consequences on target gene expression.

In summary, we generated genome-wide maps of molecular co-accessibility in the *Drosophila* genome. We showed that CRLs contain enhancers and promoters with coordinated chromatin opening and that these maps identify functional dependencies between them.

**Fig. 7.**
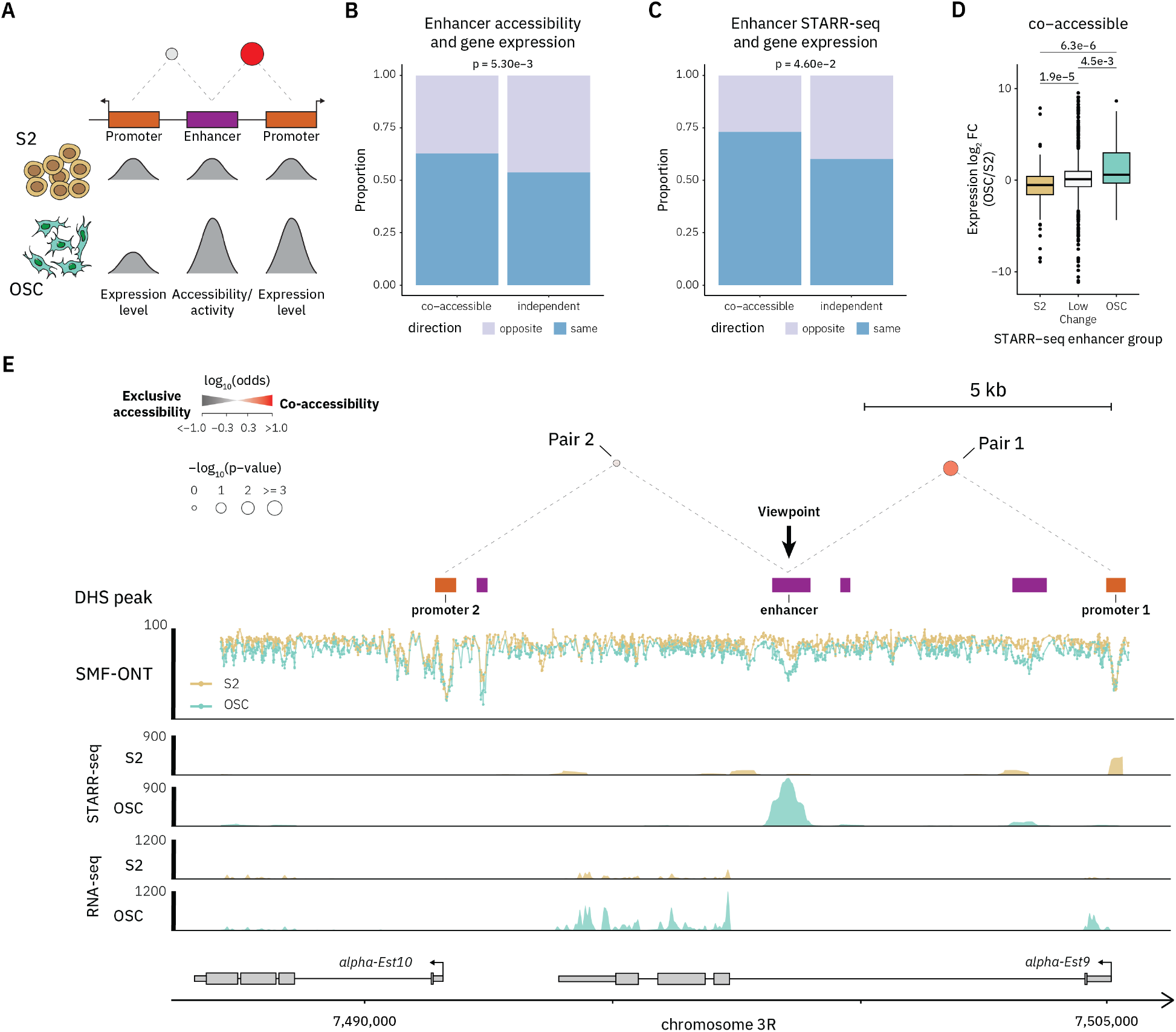
Co-accessibility identifies functional dependencies between enhancers and their target promoter. **(A)** Schematic representation of the hypothesis tested. If co-accessibility identifies dependencies between enhancers and promoters then changes in enhancer activity should lead to concordant changes in promoter activity and gene expression. **(B)** Stacked bar plot comparing the proportion of enhancer-promoter pairs for which changes in chromatin accessibility at the enhancer is correlated with concordant changes in gene expression. Statistical comparisons between groups were performed using a one-sided Fisher’s exact test. **(C)** Stacked bar plot comparing the proportion of enhancer-promoter pairs for which changes in activity as measured by STARR-seq at the enhancer is correlated with concordant changes in gene expression. Same representation as **(B). (D)** Boxplots representing the distribution of gene expression fold change (OSC vs S2) across enhancer activity groups: S2-biased (STARR-seq, *p*-value < 0.05 and fold change OSC vs S2 < −4), OSC-biased (STARR-seq, *p*-value < 0.05 and fold change OSC vs S2 > 4), and low-change (all others). The middle line of each box represents the median; boxes indicate the interquartile range (IQR; 25^th^ to 75^th^ percentile); whiskers extend to 1.5 × IQR. Statistical comparisons between groups were performed using the Wilcoxon rank-sum test. **(E)** Pair-wise map of co-accessibility illustrating how co-accessibility links enhancer to their target promoter. Example of an enhancer with increased chromatin accessibility and activity in OSC over S2 cells. This enhancer is co-accessible with the promoter of the *alpha-Est9* gene (Pair 1) that is upregulated in OSC, but not with the promoter of the *alpha-Est10* gene at the same locus. The upper panel is the same representation of Figure 2D. The lower genomic tracks show DHS peaks, SMF signal (1 *−* methylation (%)) of individual cytosines, STARR-seq, and RNA-seq signal. Tracks from S2 and OSC cells are represented in yellow and green, respectively.

## Discussion

A central question in metazoan gene regulation is to understand if the function of enhancers and promoters is coordinated to activate gene expression. Here, we tested if chromatin opening, the first step in their activation, occurs in a coordinated fashion on individual DNA molecules. At each CRL, we identified specific subsets of enhancers and promoters where chromatin opening is coordinated. We show that the *cis*-regulatory activity of these enhancers is correlated across cell types, suggesting a functional relation between them. Moreover, we connect the activity of these enhancers to changes in transcription at their target gene.

Co-accessibility suggests that each CRE may be connected to a specific set of CREs in *cis*. We observed coaccessibility for CREs located up to 30 kb apart, compatible with previous knowledge of enhancer-promoter interactions in *Drosophila* (3, 26). Yet, this is likely to be an underestimate as the current dataset cannot resolve pairs beyond 30 kb. We anticipate that increasing read length and sequencing depth may reveal interactions across longer genomic distances in the future. Co-accessibility is not restricted to enhancer-promoter pairs, and that many enhancer-enhancer and promoter-promoter pairs are co-accessible. This is consistent with reports of co-occupancy of RNA Polymerase II at neighbouring promoters (20) and the observation of coordinated transcription of functionally related genes (18, 19). Integration of the co-accessibility maps across an entire CRL suggests the existence of complex multi-way interactions between enhancers and promoters. This is in line with previous reports of the existence of 3D hubs connecting a promoter with multiple enhancers (2, 19, 46). Modelling coaccessibility data may provide a framework to infer the logical dependencies between CREs of a CRL and understand mechanisms underlying enhancer hub formation.

The observation that chromatin in distant CREs is simultaneously open at a high frequency may be explained by several molecular mechanisms. For instance, coordinated activity of distant CREs could be supported by simultaneous TF binding that would form a molecular bridge through protein-protein interactions. This model seems however unlikely given the short residency time of most TFs, temporal disconnection with the dynamics of chromatin opening (47), and the slower kinetics of 3D loop formation (48, 49). In line with this, we observe that while certain homotypic pairs of insulator proteins show significant co-occupancy between distant loci, these occur at frequencies that are unlikely to fully explain co-accessibility between CREs. Alternatively, increased co-accessibility could be the result of the exposure of two CREs to the same transcriptional hubs where TFs, cofactors and Pol II are concentrated (10, 50–55). This spatial proximity could be for instance mediated by tethering elements that were recently found to be critical to activate transcription by bringing enhancers in spatial proximity to their promoters (56). This is compatible with our observation that co-accessible CREs have high chromatin contact frequency, and the enrichment of insulator proteins at co-accessible enhancers, including the GAGA-associated factor that was previously suggested to play a role in enhancer-promoter communication (44). Moreover, it aligns with previous findings that general transcription factors (GTFs), such as the Mediator complex, participate in enhancer–promoter communication (57), even in the absence of a direct physical bridge between CREs (58).

Altogether, we demonstrate that molecular coaccessibility can be used to map the functional dependencies between CREs genome-wide. The methodology is portable and we anticipate its future application to map the architecture of CRLs across multiple cellular contexts, and its future adaptation for mammalian genomes.

## Methods

### Cell culture

*Drosophila* embryonic S2 cells were cultured at 25 °C (without CO_2_) in Schneider’s *Drosophila* medium (Gibco) supplemented with 10% of inactivated fetal bovine serum. Cells were split every 3 to 5 days at 2 million cells per mL for a maximum of 20 passages. *Drosophila* ovarian OSC cells were cultured at 25 °C (without CO_2_) in Cross and Sang’s M3 medium (Shields and Sang M3 Insect Medium supplemented with 0.6 mg/ml glutathione [Sigma, G6013], 10 mU/ml insulin [Sigma, I18821], 10% inactivated fetal bovine serum and 5% fly extract 20X from VDRC). Cells were split once/twice a week diluted 1:5 for no more than 20 passages. For splitting, OSCs were washed in PBS and trypsinized (with trypsin-EDTA) for 2 min at 25 °C.

### Expression and purification of M.CviPI

The pBAD_HisMBP3C-M.CviPI construct was freshly transformed into *E. coli* NEBExpress^®^ Iq cells. Expression cultures were grown at 37 °C in LB medium supplemented bioR*χ*iv | 11 with 0.2% glucose and 100 µg/mL Carbenicillin until OD600 0.8. Temperature was reduced to 18 °C and 0.02% L-arabinose was added to the cultures to induce M.CviPI expression. After overnight expression at 18 °C, the cultures were harvested by centrifugation. The cell pellets were resuspended in lysis buffer (50 mM Tris pH 8.0, 1 M NaCl, 10% glycerol, 0.01 mg/mL DNase, 2 mM MgCl_2_ and cOmpleteTM EDTA-free protease inhibitors) and lysed using a microfluidizer device, followed by centrifugation. Recombinant M.CviPI was purified from the cleared E. coli lysate using a combination of MBPTrap and Ni-NTA affinity chromatography, Heparin chromatography and anion exchange chromatography (HiTrap Q HP column). The final purified M.CviPI protein was stored at −20 °C in 25 mM Tris pH 8.0, 50 mM NaCl and 50% glycerol at a concentration of 1 mg/mL. A more detailed expression and purification protocol can be found at protocols.io (59).

### SMF-ONT

Single-molecule footprinting protocol was adapted from Kleinendorst and Barzaghi *et al*., 2021 (37) and optimised for long-read sequencing and high weight DNA extraction. Intact nuclei were extracted by re-suspending 2.5 *×* 10^6^ *Drosophila* cells (S2 or OSC) in ice-cold hypertonic lysis buffer (10 mM Tris pH = 7.4, 10 mM NaCl, 3 mM MgCl_2_, 0.1 mM EDTA, 0.5% IGEPAL). After 10 min incubation on ice, nuclei were spun down and washed with a lysis buffer (10 mM Tris pH = 7.4, 10 mM NaCl, 3 mM MgCl_2_, 0.1 mM EDTA). Nuclei were spun down and resuspended, using wild-bore tips, in M.CviPI reaction buffer (50 mM Tris pH 8.5, 50 mM NaCl, 10 mM DTT) supplemented with 0.6 mM SAM and 300 mM Sucrose. The nuclei were then incubated with 200U of M.CviPI (EMBL PEPCORE) at 30 °C for 7.5 min. A second incubation round was performed with 100U of M.CviPI and 128 pmol of SAM (NEB, B9003S) at 30 °C for 7.5 min. A final incubation to methylate accessible CpG di-nucleotides was performed with 60U of M.SssI (NEBM0226L), 10 mM MgCl_2_ and 128 pmol of SAM at 30 °C for 7.5 min. At each step, reaction tubes were mixed by inversion or flicking to avoid DNA shearing. Methyltransferases were heat-deactivated with a 20 min incubation at 65 °C. After an overnight Proteinase K (200 µg/mL) digestion at 55 °C, high weight molecular DNA was extracted using the Quick-DNA HMW MagBead kit (Zymo, D6060) following the manufacturer protocol and using wide-bore tips. DNA concentration was measured using Qubit high sensitivity dsDNA kit (ThermoFisher, Q32854) and the DNA fragment size measured using FemtoPulse 165 kb extended kit (Agilent, FP-1002-0275) following manufacturer instructions. When possible, DNA fragments > 25 kb were enriched using the Short Read Eliminator (PacBio, 102-208-300) following manufacturer protocol. Ligation library kits (ONT, SQK-LSK109 / SQK-LSK114) were used with 1 µg of input DNA to generate the sequencing libraries. Nanopore chemistries varied between samples based on availabilities at the time of processing (Table S1). MinION flow cells (R9.4.1) were loaded with 50 fmol of library while the PromthION flow cells (R10.4.1) with 20 fmol. Depending on the DNA fragment length and the DNA concentration, up to 3 reloads were performed per sequencing run using the flow cell wash kit (ONT, EXP-WSH004).

### Generation of training data for methylation calling

gDNA was extracted from *Drosophila* S2 cells using the Quick-DNA HMW MagBead kit (Zymo, D6060). gDNA fragment sizes were homogenised to 6 kb using g-Tubes (Covaris, 520079) following the manufacturer protocol. Fully methylated CpG and GpC gDNA was produced by two consecutive 30 min incubation at 37 °C with 8U/DNA µg of M.CviPI GpC methyltransferase (in house) with 1.2 mM SAM (NEB, B9003S) and one incubation with M.SssI CpG methyltransferase (NEB, M0226L) supplemented with 7.5 µM MgCl_2_ and 0.8 mM SAM. The enzymatic reaction was chemically stopped (20 mM Tris, 600 mM NaCl, 1% SDS 10 mM EDTA) with 14 µM proteinase K. After 4 hrs incubation at 55 °C, the methylated DNA was purified with phenol/chloroform and glycogen/salt DNA precipitation. The unmethylated gDNA and fully methylated DNA libraries were generated using the V14 multiplexing Nanopore ligation kit (ONT, SQK-NBD114.24).

### Training of the custom GpC, CpG methylation– caller

For each sample, Nanopore raw data were base called using dorado basecaller 0.3.4 using the base call model dna_r10.4.1_e8.1_400bps_sup@4.2.0 and mapped on the `dm6` *Drosophila* genome. All reads with a q-score below 7 were removed from the data. The methylation call model was trained using remora 3.0.0 software in a convolutional neural network framework. In summary, the training datasets was generated using the raw signal (`.pod5` file) and the base called data (`BAM` file) with the following syntax:

~~~
remora dataset prepare <pod5> <BAM> \
--refine-kmer-level-table data/ONT/9 mer_levels_v1.txt \
--refine-rough-rescale \
--motif CG 0 \
--motif GC 1 \
--max-chunks-per-read 20 \
--mod-base-control # for unmethylated sample
# or
--mod-base m 5mCG_5mGC # for fully
methylated sample
~~~

This generated 24,193,388 chunks from 1,242,986 reads for the unmethylated sample and 19,090,309 chunks from 983,748 reads for the fully methylated one. The training dataset was created by merging the two ground truth sample datasets with equivalent weight. The model was trained in 31 consecutive epoch using the command:

~~~
remora model train data/prepdata/
train_dataset_CpG_GpC.jsn \
--model data/ONT/ConvLSTM_w_ref.py \
--chunk-context 50 50
~~~

To test the model accuracy, cytosine methylation status in CpG and GpC contexts was inferred from the training data itself or another set of samples unmethylated or fully methylated `infer from_pod5_and_bam` from remora. The confusion matrix was computed using the `remora validate from_modbams` tool. The confusion matrix graph of the model (Figure S1A) was generated using the computed confusion matrix and a custom R (`v4.2.2`) script `plot_confusion_matrix.r`. Our custom methylation calling 5 kHz model corrects systematic errors observed in models trained solely on all-C methylation provided by ONT (Figures S1E-F).

### SMF-ONT Methylation calling

For the R10.4.1 4 kHz dataset (S2 cell replicate 1), methylation calling was performed using the all-C model (`v4.0.1_5mC`), available in the ONT rerio repository https://github.com/nanoporetech/rerio/tree/master/dorado_models. Comparison of the methylation output from this model with equivalent bisulfite-based SMF data from the same sample revealed no discrepancies (data not shown). For the R10.4.1 5 kHz and R9.4.1 datasets, methylation models trained on GpC and GpC contexts were used (see *Methods – Training of the custom GpC, CpG methylation-caller*) Basecalling, methylation calling, and genomic alignment were performed using Bonito `v0.7.3` (with integrated Minimap2 `v2.24`). The super accuracy model `v4.0.1` was used for 4 kHz data, while `v4.2.0` was applied to 5 kHz datasets. We used Bonito since it’s the only basecaller that supports methylation calling for cytosines in two specific dinucleotide contexts (CpG and GpC). Methylation signals were extracted from BAM files, stored in the `MM` and `ML` tags, with `pysam`, and converted into `mBED` format following https://github.com/timplab/nanopore-methylation-utilities. The resulting `mBED` files were indexed using `tabix` (HTSlib `v1.19.2`). Methylation probabilities were transformed into log-likelihood ratios (LLRs) for compatibility with legacy methylation callers. A stringent absolute LLR cutoff of 2 (corresponding to a *p*-value of approximately 0.12) was used to classify cytosines as methylated or unmethylated. Cytosines for which the model returned uncertain predictions (−2 *>* LLR *<* 2) were excluded from downstream analyses. A summary of all SMF-ONT datasets, including the basecalling and methylation models used, is provided in Table S1.

### Definition of CREs

*Cis*-regulatory elements (CREs) were defined based on DNase I hypersensitive sites (DHSs) (36, 42). Overlapping DHSs with peak summits located within 150 bp of each other were merged into a single region, resulting in 20,595 unique CREs. The median CRE size is 428 bp. We classified these CREs based on their proximity to transcriptional start sites (TSSs): those within 500 bp as proximal and those farther than 500 bp as distal CREs.

### Single molecule sorting

We adapted the principles of single-molecule analysis from Krebs *et al*., 2017 (41) and Sönmezer *et al*., 2021 (30) to accommodate the `mBED` format. For CRE accessibility, we applied a methylation-based single-molecule classifier in which methylation levels were extracted from a 100 bp window centred on each DHS summit. For each read, methylation within this window was binarized, classifying individual molecules into discrete accessibility states: “0” for accessible CREs and “1” for closed CREs.

For TF binding sites, we used a molecular classifier previously developed to quantify TF binding at single-molecule resolution (30, 41). This method extracts methylation from three bins positioned relative to the TF motif: upstream [-35:-25], motif-centred [-7:7], and downstream [25:35], designed to distinguish TF footprints from nucleosome occupancy. Methylation in each bin was binarized, generating a 3-bit vector per molecule corresponding to one of eight possible accessibility states. This enables classification into distinct patterns such as TF bound (101), fully accessible (111), or nucleosome-occluded configurations. To allow sorting of TFBS separated by less than 80 bp, the TF cluster strategy was used as described in Sönmezer *et al*., 2021 (30).

### DHS co-accessibility and TF co-binding testing

To test for the co-occurrence of chromatin accessibility or TF binding, we selected all CREs and TF binding sites that were contained within a 40 kb window. A minimum distance of 100 bp between DHSs and 15 bp between TFs was enforced to avoid overlapping sorted bins. Additionally, only features with at least 5% accessibility (for DHSs) or 5% binding frequency (for TFs) were retained for further analysis, resulting in 187,459 DHS pairs. To reduce computational load, only TF motifs highlighted in Figure 4C were included: Su(Hw), pnr, BEAF-32, Dref, msl-1, Clamp, Trl, M1BP, and ara/caup. This filtering yielded 44,281 TF motif pairs.

For each feature pair, only reads spanning all relevant bins on both sides of the pair were retained. The bin status for each read was aggregated across both anchors, resulting in four possible categories for DHSs:

- “11”: `DHS1_accessible_DHS2_accessible`
- “10”: `DHS1_accessible_DHS2_closed`
- “01”:`DHS1_closed_DHS2_accessible`
- “00”: `DHS1_closed_DHS2_closed` and 9 categories for TFs
- “101–101”: `TF1_bound_TF2_bound`
- “101–111”: `TF1_bound_TF2_unbound`
- “111–101”: `TF1_unbound_TF2_bound`
- “111–111”: `TF1_unbound_TF2_unbound`
- “101–nucleosome_pattern”: `TF1_bound_TF2_nucleosome`
- “nucleosome_pattern–101”: `TF1_nucleosome_TF2_bound`
- “111–nucleosome_pattern”: `TF1_unbound_TF2_nucleosome`
- “nucleosome_pattern–111”: `TF1_nucleosome_TF2_unbound`
- “nucleosome_pattern–nucleosome_pattern”: `TF1_nucleosome_TF2_nucleosome`

To determine whether the co-occurrence of accessibility or TF binding occurred more frequently than expected by chance, a two-sided Fisher’s exact test was performed for each feature pair. For each pair, reads were categorised into a 2×2 contingency table based on their binary state at both anchors (a) co-accessibility or co-binding, (b) exclusive accessibility or binding in anchor 1, (c) exclusive accessibility or binding in anchor 2 and (d) no accessibility or binding in both anchors. The odds ratio (OR) for co-occurrence was calculated using the standard formula:

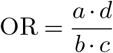

An OR of 1 indicates no association (random distribution), OR > 1 suggests positive co-occurrence (coaccessibility or co-binding), and OR < 1 suggests mutual exclusivity. The Fisher’s exact test *p*-value is computed based on the hypergeometric distribution, summing the probabilities of all possible tables with the same marginal totals that are as or more extreme than the observed one. The probability of observing a specific table is given by:

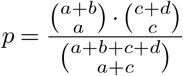

The final *p*-value is the sum of the probabilities of all contingency tables with the same marginal totals as the observed table that are as or more extreme, assuming the null hypothesis that the two features are independent (i.e., the true odds ratio is equal to 1). All odds ratios and *p*-values, along with associated metadata (e.g., genomic coordinates, distances), were compiled into summary tables for downstream interpretation (see in https://git.embl.de/grp-krebs/boulanger_2025_smf-ont).

### Proportion of co-accessible pairs as a function of coverage

To evaluate the impact of sequencing depth on the detection of co-accessible CRE pairs, we implemented a downsampling-based approach. Only CRE pairs covered by > 1,000 reads were included in the analysis (4,561 pairs in S2 and 18,812 pairs in OSC). For each pair, reads were downsampled to a fixed coverage (100–1,000 reads), and coaccessibility was then reassessed using Fisher’s exact test. The proportion of pairs showing significant co-accessibility (*p*-value < 0.05 and odds ratio > 1.5) was then calculated. This downsampling procedure was repeated 100 times, the median proportion of co-accessible pairs at each coverage level is plotted, with error bars representing 95% confidence interval (Figure S2K). We observed that the proportion of coaccessible pairs plateaus at coverage levels > 500 reads (Figure S2K). Based on this, the proportion of co-accessible pairs by pair type and distance group were calculated only for CRE pairs covered by > 500 molecules (Figure 3A-B, Figure S3AB).

### Transcription factor binding motif analysis

Transcription factor (TF) binding motifs across the `dm6` *Drosophila* genome were obtained from JASPAR 2022 database (60) and CIS-BP database (61). In total, 148 TF binding motif PWMs were scanned across the `dm6` genome with a min score of 10 using Biostring R package (`matchPWM`) (62). For each CRE, we generated a table containing TF binding motifs located within *±* 100 bp of the CRE summit. For each motif pair, we calculated the proportion of motif co-presence among CRE pairs, defined as the number of CRE pairs in which one element contained the first motif and the other element contained the second motif (regardless of order), divided by the number of pairs in which at least one element contained either motif (Figure 4A). This proportion was then compared between co-accessible CRE pairs and a distance-matched set of independent CRE pairs. The distance-matched sampling of independent CRE pairs and the comparison with co-accessible pairs were repeated 100 times. For each motif pair, we plotted the median difference in the proportion of pairs with motif co-presence across the 100 comparisons against the proportion of samplings in which co-accessible pairs exhibited a higher co-presence than distance-matched independent pairs (Figure 4B, Figure S4E). The analysis was also performed separately for each CRE pair identity (Figure S4B-D).

For each enriched motif pairs, we calculated the distribution of CRE pair identities (promoter-promoter, enhancer-promoter, and enhancer-enhancer) in which the motif pair was present. The resulting proportions were visualised as a heatmap using pheatmap R package (63) (Figure 4C, Figure 4A, 4F-G).

For the top enriched TF motif pairs, we performed TFTF co-occupancy analysis as described above. We selected TF motif pairs located within co-accessible regions as well as distance-matched independent pairs. The proportion of TFTF pairs exhibiting significant co-occupancy was then calculated and plotted (Figure 4D). We also compared the results of all Su(Hw)-Su(Hw) co-occupancy analysis that could be tested, separated by distance (Figure 4F).

### Short-read sequencing data pre-processing

A list of publicly available short-read sequencing data used in this study is shown in the Table S2. SMF data was processed as previously described (30, 37). Briefly, raw paired-end sequencing reads were trimmed using Trim Galore (`v0.6.7`) (64). The trimmed reads were aligned against a bisulfite index of the *Drosophila melanogaster* (`dm6`) genome using the R package QuasR (`v1.28.0`) (65), which uses Bowtie (66) as an aligner, with specific alignment parameters (`alignmentParameter = -e 70 -X 1000 -k` `2 -best -strata`) and keeping only uniquely aligned reads. Duplicated reads were subsequently removed using Picard (`v2.15.0`) (67). Other publicly available datasets (DNase-seq, MNase-seq, ChIP-seq, STARR-seq, and RNA-seq) were trimmed using Trim-Galore. For RNAseq, trimmed reads were aligned using STAR (`v2.7.11a`) with default parameters (68). For STARR-seq, trimmed reads were aligned using QuasR with similar parameters to the original article (`-v 3 -m 1 -best -strata -X 2000`) (42). For DNase-seq, MNase-seq, and ChIP-seq, trimmed reads were aligned using QuasR with default alignment parameters. Duplicated reads were subsequently removed using Picard (`v2.15.0`) (67).

### Micro-C data analysis

Publicly available Micro-C data from S2 cells was obtained in `Cooler` file format from GEO GSE246517 (45). The original 200-bp resolution data were coarsened to 600-bp resolution to reduce data sparsity. The resulting file was normalised using iterative correction via `cooler balance` from the Cooler tool (`v0.10.3`). The balanced micro-C file was subsequently imported into R and analysed using the HiCExperiment (`v1.6.0`), GenomicInteractions (`v1.38.0`), and InteractionSet (`v1.32.0`) packages. To compare micro-C scores between co-accessible and independent CRE pairs, we first subset the interaction set to retain only those interactions overlapping CREs (*±* 100 bp of the CRE summit) at both anchors. In total, 62,450 out of 67,934 CRE pairs were retained for the analysis, consisting of 12,412 co-accessible and 50,038 independent pairs. Interactions were categorised into distance bins (1–3 kb, 3–5 kb, 5–10 kb, and > 10 kb). Balanced micro-C scores were logtransformed, and comparisons between co-accessible and independent pairs were performed within each distance bin using Wilcoxon rank-sum tests (Figure 5A, Figure S5A).

### RNA-seq data analysis

RNA-seq data from S2 and OSC cells were pre-processed as described above. Then, genelevel count table was generated using the `featureCounts` function from the Rsubread package (`v2.18.0`) (69), with gene annotations from FlyBase (release r6.62) (70). Genes with zero raw counts across all four samples were filtered out. Differential gene expression analysis was then performed using DESeq2 (`v1.44.0`) (71). Genes with BenjaminiHochberg adjusted *p*-value < 0.05 and absolute fold change (|FC|) > 1.5 were considered differentially expressed between the two cell types.

### STARR-seq data analysis

STARR-seq data from S2 and OSC cells were pre-processed as described above. Then, STARR-seq signal at each CRE (*±* 100 bp around the summit) was quantified using the `qCount` function from the QuasR package (`v1.28.0`). CREs were retained for analysis if they showed more than one count in at least one replicate from both S2 and OSC cells. Differential analysis was performed using DESeq2 (`v1.44.0`). Input libraries were included in the analysis design, using the model:`~ condition + type + condition:type`, where `condition` represents the cell type (S2 or OSC) and `type` distinguishes input and STARR-seq samples. CREs with *p*value < 0.05 and absolute fold change (|FC|) > 4 were considered to show differential activity between the two cell types.

### Distance-matched sampling procedure

To account for potential bias coming from differences in the genomic distance distributions between co-accessible and independent CRE pairs (Figure S6A-B), we employed a distance-matched subsampling approach. All CRE pairs were first binned by their genomic distances using 500 bp intervals. For each distance bin, we quantified the number of co-accessible pairs and randomly sampled an equal number of independent pairs without replacement. This strategy ensured that the distance distribution of the independent pairs matched that of the coaccessible pairs. This sampling approach was applied in analyses presented in Figure 4, Figure 6, and Figure 7.

### Co-variation of accessibility across cell types

Only CRE pairs tested in both S2 and OSC cells were retained for this analysis. CRE pairs were classified as either co-accessible or independent using more stringent criteria. Specifically, co-accessible pairs were defined as those with a *p*-value < 0.01 and odds ratio > 2, while independent pairs were defined as those with a *p*-value > 0.05. If a CRE pair was co-accessible in at least one cell type, it was classified as co-accessible. In contrast, independent pairs were those classified as independent in both cell types. In total, 7,210 co-accessible and 40,525 independent pairs were included in the analysis.

For each CRE in a pair, we computed the change in accessibility between cell types (delta accessibility; OSC S2). To control for distance-dependent effects, independent pairs were sampled to match distance distribution of co-accessible pairs, as described above. Pearson correlation was used to assess the correlation in delta accessibility between the two CREs in distance-matched independent and co-accessible pairs. This analysis was performed separately for enhancer-enhancer (Figure 6B-C), enhancer-promoter (Figure 6E-F), and promoter-promoter pairs (Figure S6G-H). Distance-matched samplings and correlation analysis for independent pairs were repeated 1,000 times (Figure S6C, S6E, S6I).

Next, we categorised each CRE based on delta accessibility (*±* 20%) into directional groups: increased accessibility in S2, increased accessibility in OSC, or low change. For CRE pairs in which both elements exhibit changes in accessibility, we calculated the proportion of pairs showing changes in the same versus opposite directions. A one-sided Fisher’s exact test was used to compare the proportion of coordinated changes between co-accessible and distance-matched independent pairs, under the hypothesis that co-accessible pairs would exhibit a higher proportion of concordant (same direction) accessibility changes than independent pairs (Figure 6D, 6G, Figure S6J). Distance-matched samplings and Fisher’s test for independent pairs were repeated 1,000 times (Figure S6D, S6F, S6K).

### Enhancer activity and gene expression analysis

For the analysis of enhancer accessibility and gene expression (Figure 7B), enhancer-promoter pairs were retained if the promoter was associated with a gene that showed RNA-seq expression in both S2 cells and OSC. Similar to the cobioR*χ*iv | 15 variation analysis, co-accessible pairs were defined as those with a *p*-value < 0.01 and odds ratio > 2 from the coaccessibility analysis, while independent pairs were defined as those with a *p*-value > 0.05. If a pair was co-accessible in at least one cell type, it was classified as co-accessible. In contrast, independent pairs were those classified as independent in both cell types. In total, 2,419 co-accessible and 17,051 independent enhancer-promoter pairs were included in the analysis. Each enhancer-promoter pair was annotated with the direction of change in enhancer accessibility (delta accessibility > 20 for OSC, delta accessibility < −20 for S2) and gene expression (RNA-seq analysis, up-regulated in S2 or OSC). The proportion of enhancer-promoter pairs showing concordant directionality (i.e., enhancer and gene expression changes in the same cell type) was then compared between co-accessible and distance-matched independent groups. Statistical significance was assessed using a onesided Fisher’s exact test. Distance-matched samplings for independent pairs were repeated 1,000 times (Figure S7A).

Next, to investigate the relationship between enhancer activity and gene expression (Figure 7C), enhancer-promoter pairs were filtered further to retain only pairs in which the enhancer contained STARR-seq. In total, 1,400 co-accessible and 9,802 independent enhancer-promoter pairs were included in the analysis. Each enhancer-promoter pair was annotated with the direction of change in enhancer activity (STARR-seq analysis, increased activity in S2 or OSC) and gene expression (RNA-seq analysis, up-regulated in S2 or OSC). The proportion of enhancer-promoter pairs showing concordant directionality was then compared between coaccessible and distance-matched independent groups. Statistical significance was assessed using a one-sided Fisher’s exact test. Distance-matched samplings for independent pairs were repeated 1,000 times (Figure S7B). In addition, coaccessible Enhancer–Promoter pairs were grouped based on changes in enhancer activity (STARR-seq analysis). Gene expression changes (log_2_ fold change between OSC and S2) for the associated genes were compared across these enhancer groups using Wilcoxon rank-sum tests (Figure 7D).

## Data availability

SMF-ONT data were deposited array express (E-MTAB-14663). The Nextflow pipeline to analyse long-read SMF is available at: https://git.embl.de/grp-krebs/nf-smfont. Analysis code, test datasets, and documentation can be found at: https://git.embl.de/grp-krebs/boulanger_2025_smf-ont.

## ACKNOWLEDGEMENTS

The authors are grateful to the members of the Krebs laboratory for helpful discussions; and Tineke Lenstra, Tim Pollex, Laura Moniot-Perron, Michela Palamin, and Colm Doyle for comments on the manuscript. The authors are grateful to EMBL GeneCore sequencing facility and DKFZ sequencing facility for their support in sequencing data generation. The authors thank the MODIS team of the EMBL Data Science Centre for their outstanding support in the development of the nanopore data management in their Lab Integrated Data platform and setting up a performant analysis framework. The authors also appreciate Jacob Scheurich and Nikolay Dobrev for advice and GpC methyltransferase production. The salary of M.B. and K.C. are supported by the Deutsche Forschungsgemeinschaft (KR 5247/1-3). The salary of R.S. was supported by the Helmholtz Association under the joint research school “HIDSS4Health Helmholtz Information and Data Science School for Health”. Research in the laboratory of A.R.K. is supported by core funding from the EMBL, Deutsche Forschungsgemeinschaft (KR 5247/1-1, KR 5247/1-2, KR 5247/1-3), and the ERC (TFCoop-101125530).

## AUTHOR CONTRIBUTIONS

M.B and A.R.K designed the study. M.B, K.C and A.R.K wrote the manuscript. M.B performed the experiments. M.B and K.C, analysed the data. R.S and M.J.B supported initial improvements and benchmarking of the ONT methyl-callers. K.L and K.R established the M.CviPI purification protocol. A.R.K supervised conduction of the experiments and the data analysis. All authors discussed the results and commented on the manuscript.

## Supplementary Figures

**Fig. S1.**
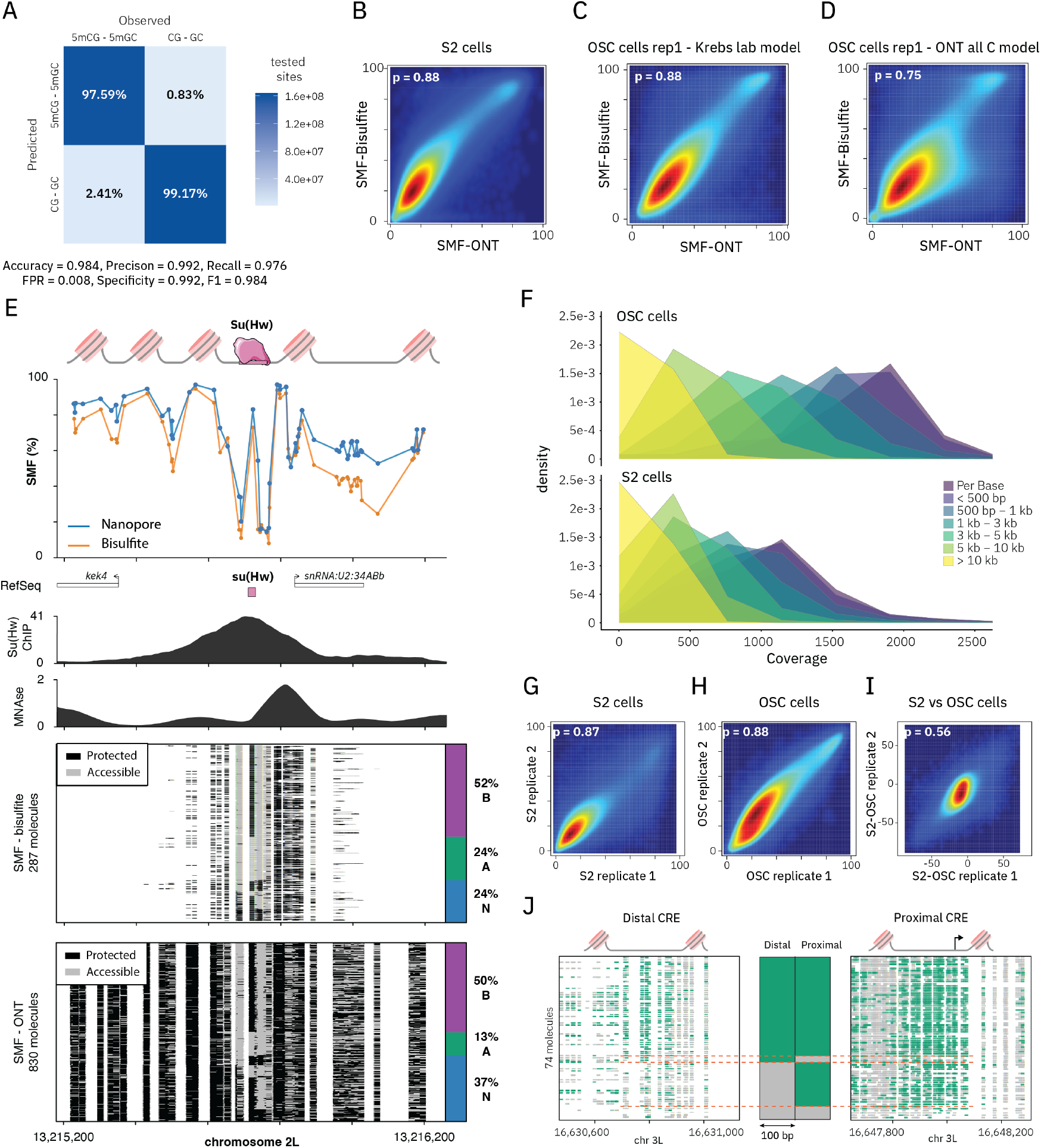
**(A)** Confusion matrix of the custom convolutional neural network model optimised for SMF-ONT data. The model, trained on fully methylated (CpG and GpC dinucleotides only) and fully unmethylated DNA prepared with LSK-114 ONT kit, detects methylation with 98.4% accuracy on 5 kHz ONT data. **(B)** SMF-ONT recapitulates reference SMF data generated by bisulfite sequencing. Density scatter plot showing the correlation of cytosine methylation (CpG and GpC contexts) between bisulfite and ONT sequencing in S2 cells (Pearson correlation: 0.88). **(C)** SMF-ONT recapitulates bisulfite-sequenced SMF data in OSC cells. Same representation as **(B). (D)** Density scatter plot showing a lower correlation between cytosine methylation measured by bisulfite sequencing and ONT sequencing using the default ONT all-cytosine methylation calling model. Same representation as **(B). (E)** Single-locus example illustrating the reproducible footprint detection of SMF-ONT using bisulfite SMF data as a reference. The upper panel displays average SMF-ONT (blue) and bisulfite (yellow) signals (1 *−* methylation (%)). Middle panels show ChIP-seq signals for Su(Hw) and MNase-seq nucleosome occupancy. Lower panels display single DNA molecule stacks from bisulfite (upper) and Nanopore (lower) sequencing across a 1,000 bp window around the Su(Hw) binding site, with cytosine methylation status colour-coded (black: protected from methylation; grey: accessible) identical as Figure 4E. Stacked bar charts represent the fraction of molecules bound to the transcription factor (TF bound, purple), with accessible motifs (green) and nucleosome-bound regions (blue). **(F)** Density plots showing coverage as a function of genomic distance between CRE pairs in OSC (upper panel) and S2 (lower panel) datasets. **(G-H)** Density scatter plots showing reproducibility of SMF-ONT cytosine methylation data between biological replicates in **(H)** S2 cells (Pearson correlation: 0.87) and **(H)** OSC cells (Pearson correlation: 0.88). **(I)** Density scatter plots showing correlation of differential cytosine methylation between S2 and OSC cells across two replicates (Pearson correlation: 0.56). **(J)** Zoomed-in view illustrating co-accessibility between the promoter of the *Abl* gene and a putative enhancer located 17.2 kb downstream. Left and right panels display the same individual SMF-ONT reads over the two regions, same colour code as in Figure 1C. The middle panel summarises the accessibility status of each CRE (grey: protected from methylation; green: accessible).

**Fig. S2.**
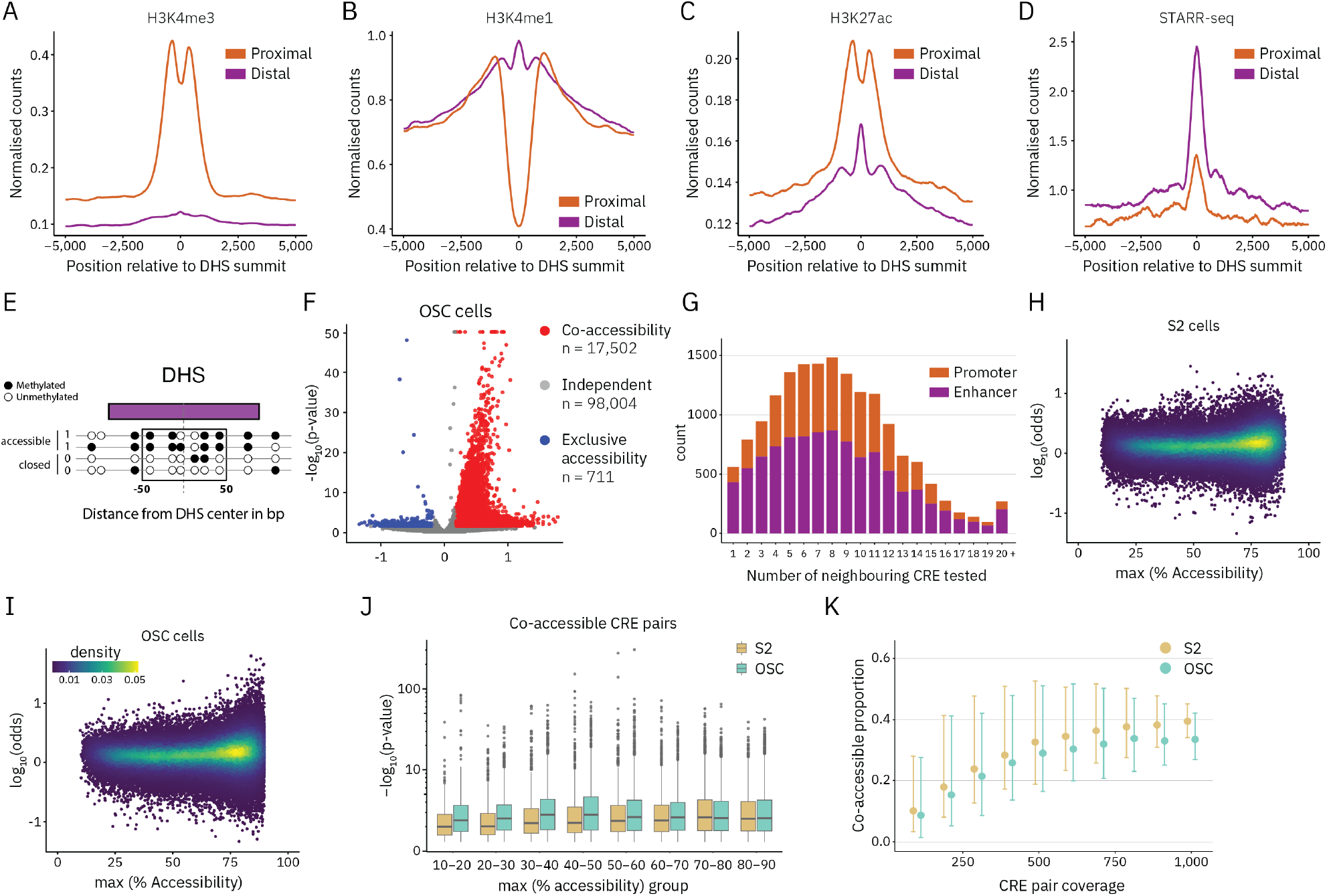
**(A-D)** Composite plot showing the average signal for proximal (orange) and distal (purple) CREs within a 10 kb window centred on the DHS summit for: H3K4me3 ChIP-seq **(A)**, H3K4me1 ChIP-seq **(B)**, H3K27ac ChIP-seq **(C)**, and STARR-seq **(D). (E)** Schematic representation of cytosine methylation quantification at DNase I hypersensitive sites (DHSs). Cytosines are depicted as circles: black indicates methylated cytosines; white indicating unmethylated ones. For each single DNA molecule, the average cytosine methylation level is calculated within a 100 bp window centred at the DHS summit. Based on their average methylation profile within the window, each read is then classified into accessible (*≥* 50% methylation) and closed (< 50% methylation). **(F)** Volcano plot illustrating the results of co-accessibility tests in OSC cells. Colours represent the result of the test (red: co-accessibility; blue: exclusive accessibility; grey: independent). **(G)** Histogram showing the distribution of the number of neighbouring CREs tested for co-accessibility per CRE. Bar colours indicate CRE type based on proximity to a transcription start site (TSS): DHSs within 500 bp of a TSS are classified as promoters (orange), while more distal DHSs are considered enhancers (purple). **(H-I)** Density scatter plot showing log(odds) from the Fisher’s exact test as a function of the highest accessibility level of the two CREs in a pair being tested in S2 cells **(H)** and OSC **(I). (J)** Box plot showing the *p*-value from the Fisher’s exact test as a function of the highest accessibility level of the two CREs in a pair being tested in S2 cells and OSC. **(K)** Proportion of co-accessible CRE pairs as a function of sequencing coverage in S2 and OSC cells. Only CRE pairs with > 1,000× coverage were included in the analysis (4,561 pairs in S2 and 18,812 pairs in OSC). For each pair, reads were randomly downsampled to coverage levels ranging from 1,000 to 100 reads. Co-accessibility was then assessed using Fisher’s exact test, and the proportion of pairs showing significant co-accessibility (*p*-value < 0.05 and odds ratio > 1.5) was calculated. This downsampling procedure was repeated 100 times. Error bars represent 95% confidence intervals across 100 iterations.

**Fig. S3.**
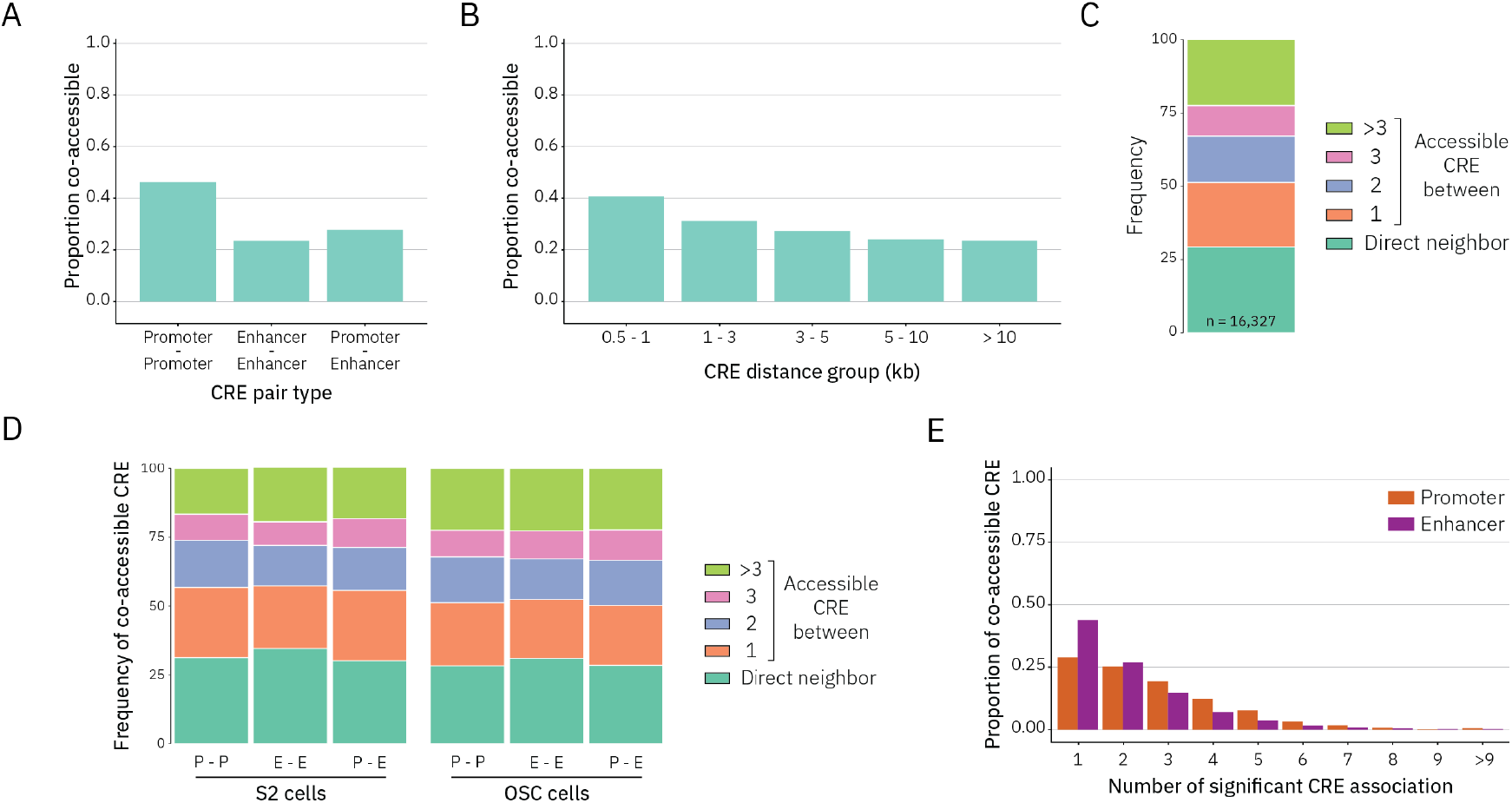
**(A)** Bar chart displaying the proportion of co-accessible CRE pairs categorised by pair type in OSC cells. **(B)** Bar charts illustrating the proportion of co-accessible CRE pairs as a function of genomic distance between them. Only pairs with > 500x coverage in OSC cells were included in this analysis to avoid false negatives due to low coverage. **(C)** Stacked bar plots depicting the number of CREs intercalated between two co-accessible CREs in OSC cells. **(D)** Stacked bar plots depicting the number of CREs located between two co-accessible CREs, separated by pair type (P-P: promoter-promoter, E-E: enhancer-enhancer, E-P: enhancer-promoter), in S2 and OSC cells. **(E)** Histogram showing the distribution of significant co-accessible pairs per CRE, separated by CRE type (promoter and enhancer).

**Fig. S4.**
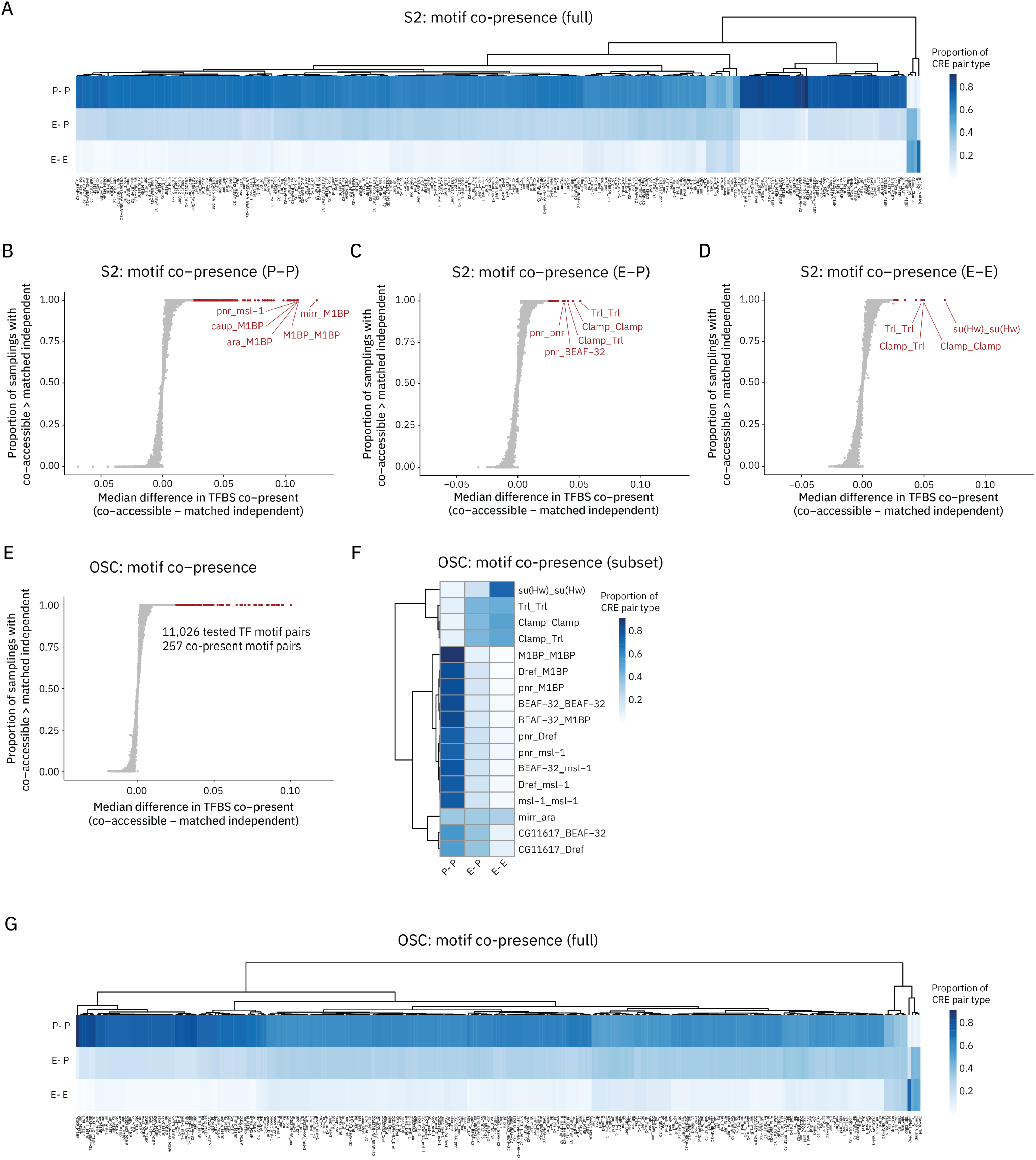
**(A)** Heatmap showing the distribution of 248 co-present TF motif pairs across different types of co-accessible CRE pairs in S2 cells. The colour scale represents the proportion of each CRE pair type (promoter-promoter, enhancer-promoter, enhancer-enhancer) in which the motif pairs are located. **(B-D)** Scatter plots displaying TF motif co-presence in co-accessible pair versus distance-matched independent pairs in S2 cells. The analysis was performed on **(B)** promoter-promoter, **(C)** enhancer-promoter, and **(D)** enhancer-enhancer pairs. The x-axis represents the median difference in the proportion of pairs with motif co-presence between co-accessible and 100 distance-matched samplings of independent pairs. The y-axis shows the proportion of samplings in which co-accessible pairs exhibit a higher motif co-presence than distance-matched independent pairs. Top candidate motif pairs with the highest median difference are highlighted in red. **(E)** Scatter plot displaying TF motif co-presence in co-accessible pair versus distance-matched independent pairs in OSC cells. Same representation as **(B). (F)** Heatmap showing the distribution of top co-present TF motif pairs across different types of co-accessible CRE pair in OSC cells. Same representation as **(A). (G)** Heatmap showing the distribution of 257 co-present TF motif pairs across different types of co-accessible CRE pair in OSC cells. Same representation as **(A)**.

**Fig. S5.**
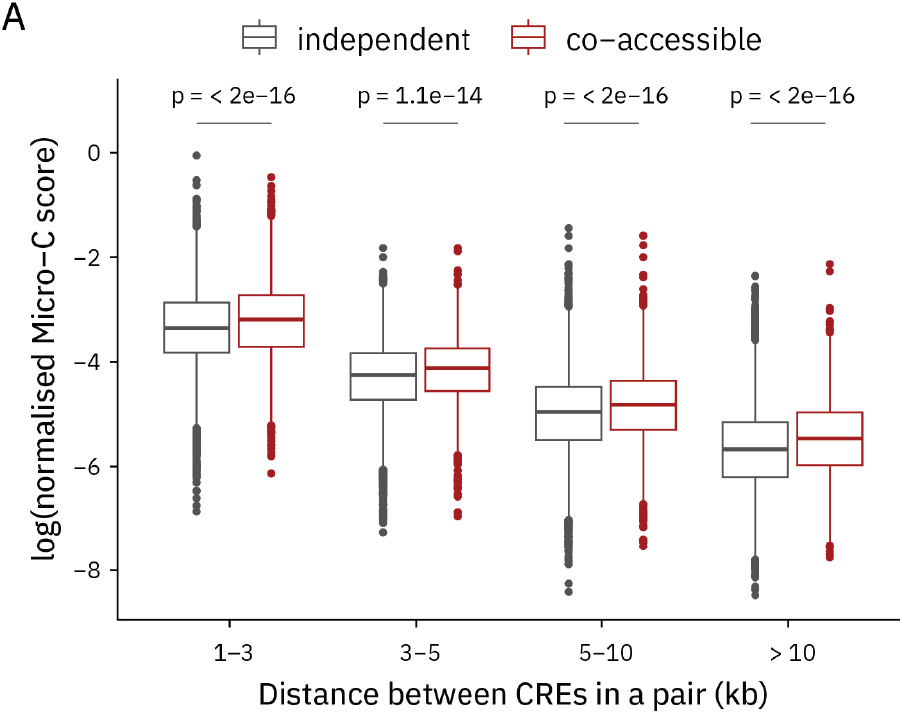
**(A)** Boxplots representing the distribution of balanced Micro-C contact frequency scores between CRE pairs, comparing independent (grey) and co-accessible (red) pairs. The analysis is stratified by the distance between pairs. The statistical comparisons between groups were performed using the Wilcoxon rank-sum test.

**Fig. S6.**
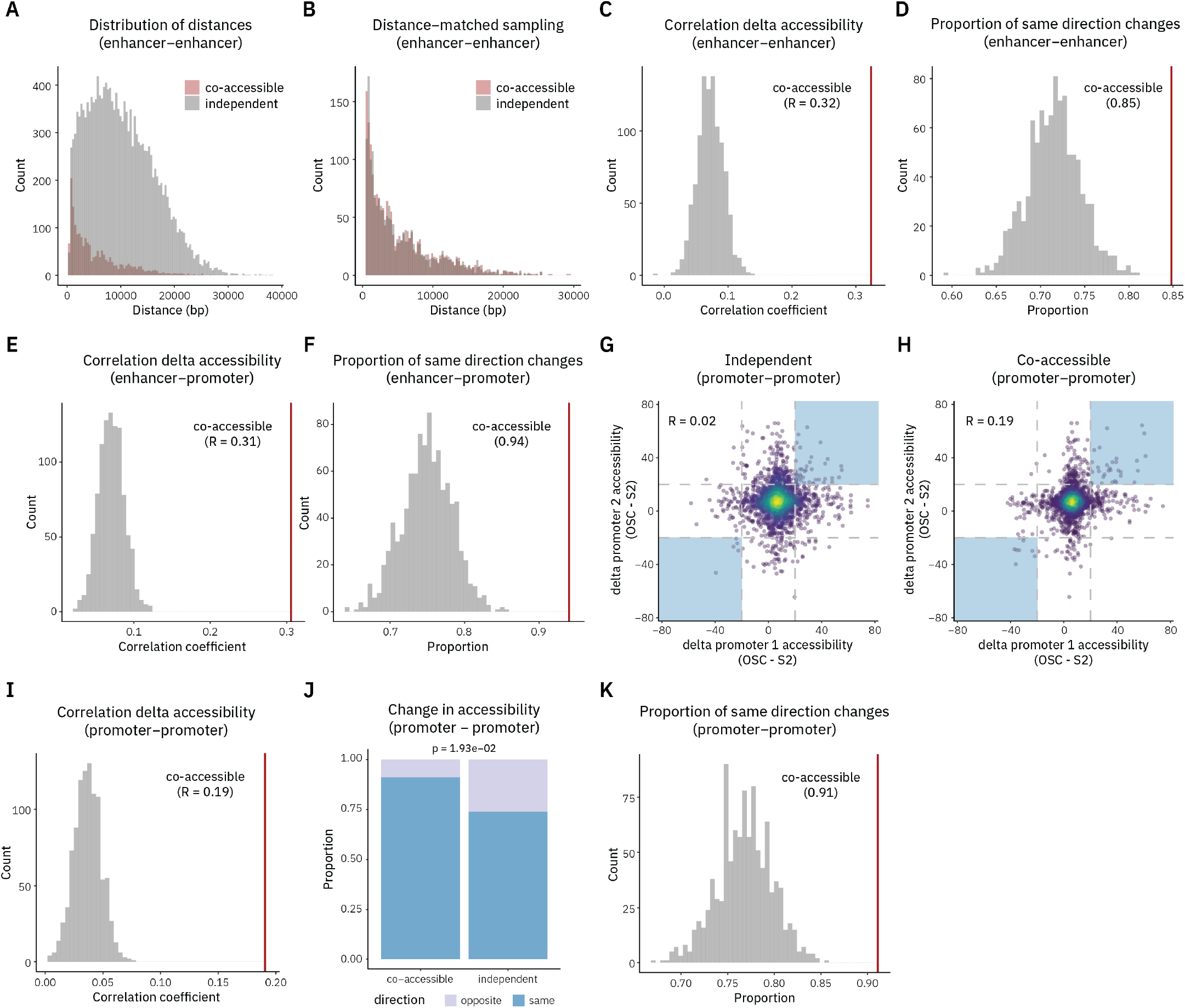
**(A)** Histogram showing genomic distances between enhancer-enhancer pairs for co-accessible (red) and independent pairs (grey). **(B)** Histogram showing genomic distances between enhancer-enhancer pairs for co-accessible (red) and a distance-matched independent pairs (grey). Independent enhancer-promoter pairs were binned by distance (500 bp bins), and an equal number of pairs were randomly sampled from each bin to match the distribution of co-accessible pairs. **(C)** Histogram showing Pearson correlation coefficients from 1,000 random samplings of distance-matched independent enhancer-enhancer pairs (grey). The observed correlation for co-accessible pairs is shown as a red line (R = 0.32). **(D)** Histogram showing the proportion of distance-matched independent enhancer-enhancer pairs with concordant changes in chromatin accessibility, across 1,000 samplings (grey). The red line marks the observed value for co-accessible pairs (0.85). **(E)** Histogram showing Pearson correlation coefficients from 1,000 random samplings of distance-matched independent enhancer-promoter pairs. Same representation as **(C). (F)** Histogram showing the proportion of distance-matched independent enhancer-promoter pairs with concordant changes in chromatin accessibility. Same representation as **(D). (G-H)** Cell type-specific chromatin accessibility changes are coordinated at co-accessible promoter-promoter pairs. Scatter plot showing the pairwise difference in chromatin accessibility between cell types at **(G)** independent and **(H)** co-accessible promoter-promoter pairs. Independent pairs were sampled to be distance-matched to the co-accessible pairs. Blue shaded boxes highlight promoter-promoter pairs exhibiting highly coordinated changes in accessibility (> 20%). Pearson correlation coefficient is displayed in the plot. **(I)** Histogram showing Pearson correlation coefficients from 1,000 random samplings of distance-matched independent promoter-promoter pairs. Same representation as **(C). (J)** Stacked bar plot comparing the proportion of promoter-promoter pairs with coordinated chromatin accessibility changes across cell types. Only pairs with high variation in the accessibility of both promoters (> 20%) were included. Statistical comparisons between groups were performed using a one-sided Fisher’s exact test. **(K)** Histogram showing the proportion of distance-matched independent promoter-promoter pairs with concordant changes in chromatin accessibility. Same representation as **(D)**.

**Fig. S7.**
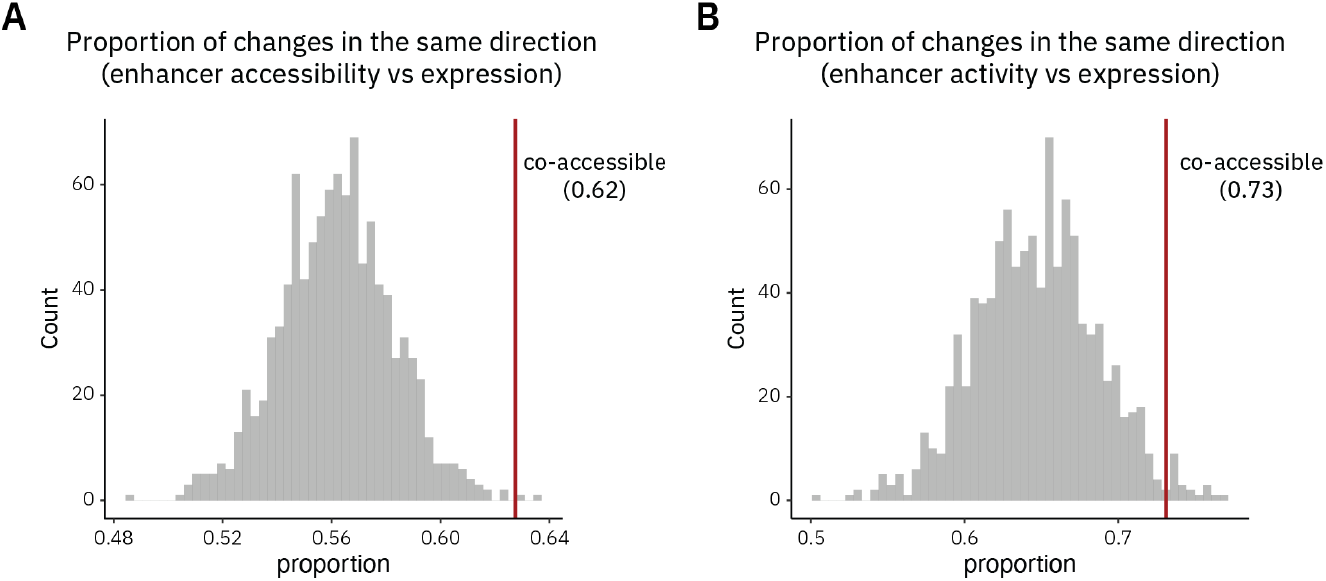
**(A)** Histogram showing the proportion of distance-matched independent enhancer–promoter pairs with concordant changes in enhancer accessibility (SMF-ONT) and gene expression (RNA-seq), across 1,000 samplings (grey). The red line marks the observed value for co-accessible pairs (0.62). **(B)** Histogram showing the proportion of distance-matched independent enhancer–promoter pairs with concordant changes in enhancer activity (STARR-seq) and gene expression (RNA-seq). Same representation as **(A)**.

## Supplementary Table

**Table S1.**
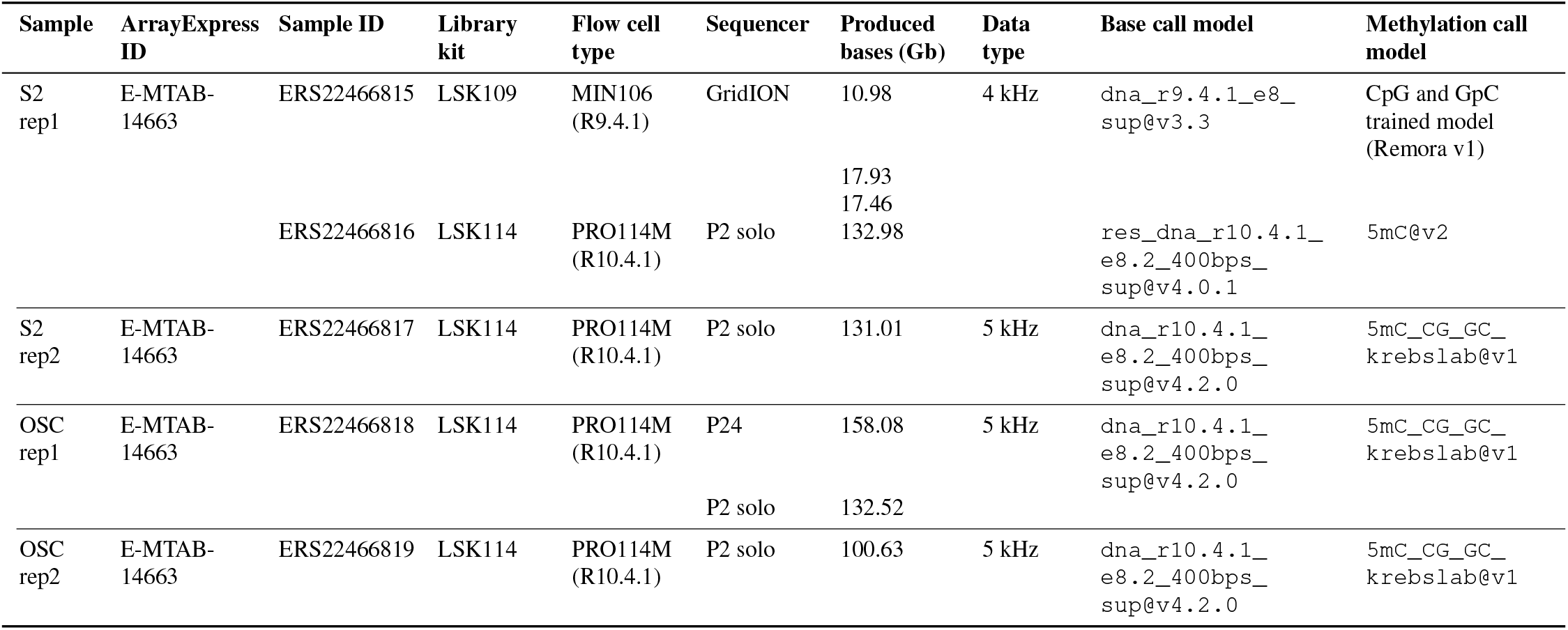
Summary of the SMF-ONT datasets.

**Table S2.**
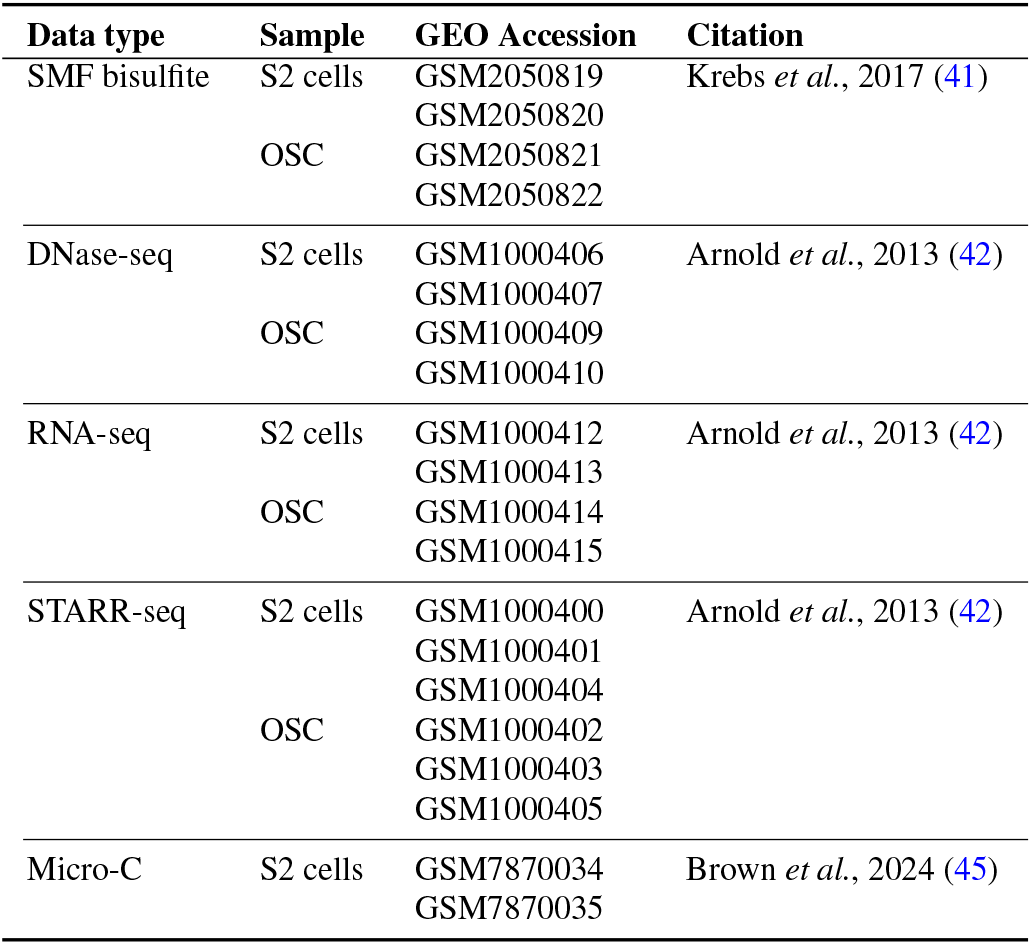
Publicly available data used.

## References

1. Andreas R. Gschwind, Kristy S. Mualim, Alireza Karbalsayghareh, Maya U. Sheth, Kushal K. Dey, Evelyn Jagoda, Ramil N. Nurtdinov, Wang Xi, Anthony S. Tan, Hank Jones, X. Rosa Ma, David Yao, Joseph Nasser, Žiga Avsec, Benjamin T. James, Muhammad S. Shamim, Neva C. Durand, Suhas S. P. Rao, Ragini Mahajan, Benjamin R. Doughty, Kalina Andreeva, Jacob C. Ulirsch, Kaili Fan, Elizabeth M. Perez, Tri C. Nguyen, David R. Kelley, Hilary K. Finucane, Jill E. Moore, Zhiping Weng, Manolis Kellis, Michael C. Bassik, Alkes L. Price, Michael A. Beer, Roderic Guigó, John A. Stamatoyannopoulos, Erez Lieberman Aiden, William J. Greenleaf, Christina S. Leslie, Lars M. Steinmetz, Anshul Kundaje, and Jesse M. Engreitz. An encyclopedia of enhancer-gene regulatory interactions in the human genome. bioRxiv: The Preprint Server for Biology, page 2023.11.09.563812, November 2023. ISSN 2692-8205. doi: 10.1101/2023.11.09.563812.

2. Christopher M. Uyehara and Effie Apostolou. 3D enhancer-promoter interactions and multiconnected hubs: Organizational principles and functional roles. Cell Reports, 42(4):112068, April 2023. ISSN 2211-1247. doi: 10.1016/j.celrep.2023.112068.

3. Eileen E. M. Furlong and Michael Levine. Developmental enhancers and chromosome topology. Science (New York, N.Y.), 361(6409):1341–1345, September 2018. ISSN 1095-9203. doi: 10.1126/science.aau0320.

4. Argyris Papantonis and A. Marieke Oudelaar. Mechanisms of Enhancer-Mediated Gene Activation in the Context of the 3D Genome. Annual Review of Genomics and Human Genetics, March 2025. ISSN 1527-8204, 1545-293X. doi: 10.1146/annurev-genom-120423-012301.

5. Jacques P. Bothma, Hernan G. Garcia, Samuel Ng, Michael W. Perry, Thomas Gregor, and Michael Levine. Enhancer additivity and non-additivity are determined by enhancer strength in the Drosophila embryo. eLife, 4:e07956, August 2015. ISSN 2050-084X. doi: 10.7554/eLife.07956.

6. Deborah Hay, Jim R. Hughes, Christian Babbs, James O. J. Davies, Bryony J. Graham, Lars Hanssen, Mira T. Kassouf, A. Marieke Marieke Oudelaar, Jacqueline A. Sharpe, Maria C. Suciu, Jelena Telenius, Ruth Williams, Christina Rode, Pik-Shan Li, Len A. Pennacchio, Jacqueline A. Sloane-Stanley, Helena Ayyub, Sue Butler, Tatjana Sauka-Spengler, Richard J. Gibbons, Andrew J. H. Smith, William G. Wood, and Douglas R. Higgs. Genetic dissection of the α-globin super-enhancer in vivo. Nature Genetics, 48(8):895–903, August 2016. ISSN 1546-1718. doi: 10.1038/ng.3605.

7. Joung-Woo Hong, David A. Hendrix, and Michael S. Levine. Shadow enhancers as a source of evolutionary novelty. Science (New York, N.Y.), 321(5894):1314, September 2008. ISSN 1095-9203. doi: 10.1126/science.1160631.

8. Marco Osterwalder, Iros Barozzi, Virginie Tissières, Yoko Fukuda-Yuzawa, Brandon J. Mannion, Sarah Y. Afzal, Elizabeth A. Lee, Yiwen Zhu, Ingrid Plajzer-Frick, Catherine S. Pickle, Momoe Kato, Tyler H. Garvin, Quan T. Pham, Anne N. Harrington, Jennifer A. Akiyama, Veena Afzal, Javier Lopez-Rios, Diane E. Dickel, Axel Visel, and Len A. Pennacchio. Enhancer redundancy provides phenotypic robustness in mammalian development. Nature, 554(7691):239–243, February 2018. ISSN 1476-4687. doi: 10.1038/nature25461.

9. Michael W. Perry, Alistair N. Boettiger, and Michael Levine. Multiple enhancers ensure precision of gap gene-expression patterns in the Drosophila embryo. Proceedings of the National Academy of Sciences of the United States of America, 108(33):13570–13575, August 2011. ISSN 1091-6490. doi: 10.1073/pnas.1109873108.

10. Denes Hnisz, Brian J. Abraham, Tong Ihn Lee, Ashley Lau, Violaine Saint-André, Alla A. Sigova, Heather A. Hoke, and Richard A. Young. Super-enhancers in the control of cell identity and disease. Cell, 155(4):934–947, November 2013. ISSN 1097-4172. doi: 10.1016/j.cell.2013.09.053.

11. Warren A. Whyte, David A. Orlando, Denes Hnisz, Brian J. Abraham, Charles Y. Lin, Michael H. Kagey, Peter B. Rahl, Tong Ihn Lee, and Richard A. Young. Master transcription factors and mediator establish super-enhancers at key cell identity genes. Cell, 153(2): 307–319, April 2013. ISSN 1097-4172. doi: 10.1016/j.cell.2013.03.035.

12. Denes Hnisz, Jurian Schuijers, Charles Y. Lin, Abraham S. Weintraub, Brian J. Abraham, Tong Ihn Lee, James E. Bradner, and Richard A. Young. Convergence of developmental and oncogenic signaling pathways at transcriptional super-enhancers. Molecular Cell, 58 (2):362–370, April 2015. ISSN 1097-4164. doi: 10.1016/j.molcel.2015.02.014.

13. Henry F. Thomas, Elena Kotova, Swathi Jayaram, Axel Pilz, Merrit Romeike, Andreas Lackner, Thomas Penz, Christoph Bock, Martin Leeb, Florian Halbritter, Joanna Wysocka, and Christa Buecker. Temporal dissection of an enhancer cluster reveals distinct temporal and functional contributions of individual elements. Molecular Cell, 81(5):969–982.e13, March 2021. ISSN 1097-4164. doi: 10.1016/j.molcel.2020.12.047.

14. Joseph W. Blayney, Helena Francis, Alexandra Rampasekova, Brendan Camellato, Leslie Mitchell, Rosa Stolper, Lucy Cornell, Christian Babbs, Jef D. Boeke, Douglas R. Higgs, and Mira Kassouf. Super-enhancers include classical enhancers and facilitators to fully activate gene expression. Cell, 186(26):5826–5839.e18, December 2023. ISSN 1097-4172. doi: 10.1016/j.cell.2023.11.030.

15. Grace Bower, Ethan W. Hollingsworth, Sandra H. Jacinto, Joshua A. Alcantara, Benjamin Clock, Kaitlyn Cao, Mandy Liu, Adam Dziulko, Ana Alcaina-Caro, Qianlan Xu, Dorota Skowronska-Krawczyk, Javier Lopez-Rios, Diane E. Dickel, Anaïs F. Bardet, Len A. Pennacchio, Axel Visel, and Evgeny Z. Kvon. Range extender mediates long-distance enhancer activity. Nature, 643(8072):830–838, July 2025. ISSN 1476-4687. doi: 10.1038/s41586-025-09221-6.

16. Henry F. Thomas, Songjie Feng, Felix Haslhofer, Marie Huber, María García Gallardo, Vincent Loubiere, Daria Vanina, Mattia Pitasi, Alexander Stark, and Christa Buecker. Enhancer cooperativity can compensate for loss of activity over large genomic distances. Molecular Cell, 85(2):362–375.e9, January 2025. ISSN 1097-4164. doi: 10.1016/j.molcel.2024.11.008.

17. Takashi Fukaya, Bomyi Lim, and Michael Levine. Enhancer Control of Transcriptional Bursting. Cell, 166(2):358–368, July 2016. ISSN 1097-4172. doi: 10.1016/j.cell.2016.05.025.

18. Michal Levo, João Raimundo, Xin Yang Bing, Zachary Sisco, Philippe J. Batut, Sergey Ryabichko, Thomas Gregor, and Michael S. Levine. Transcriptional coupling of distant regulatory genes in living embryos. Nature, 605(7911):754–760, May 2022. ISSN 1476-4687. doi: 10.1038/s41586-022-04680-7.

19. Tim Pollex, Raquel Marco-Ferreres, Lucia Ciglar, Yad Ghavi-Helm, Adam Rabinowitz, Rebecca Rodriguez Viales, Christoph Schaub, Aleksander Jankowski, Charles Girardot, and Eileen E. M. Furlong. Chromatin gene-gene loops support the cross-regulation of genes with related function. Molecular Cell, 84(5):822–838.e8, March 2024. ISSN 1097-4164. doi: 10.1016/j.molcel.2023.12.023.

20. Thomas W. Tullius, R. Stefan Isaac, Danilo Dubocanin, Jane Ranchalis, L. Stirling Churchman, and Andrew B. Stergachis. RNA polymerases reshape chromatin architecture and couple transcription on individual fibers. Molecular Cell, page S1097276524006671, August 2024. ISSN 10972765. doi: 10.1016/j.molcel.2024.08.013.

21. Dig B. Mahat, Nathaniel D. Tippens, Jorge D. Martin-Rufino, Sean K. Waterton, Jiayu Fu, Sarah E. Blatt, and Phillip A. Sharp. Single-cell nascent RNA sequencing unveils coordinated global transcription. Nature, 631(8019):216–223, July 2024. ISSN 1476-4687. doi: 10.1038/s41586-024-07517-7.

22. Seungsoo Kim and Joanna Wysocka. Deciphering the multi-scale, quantitative cisregulatory code. Molecular Cell, 83(3):373–392, February 2023. ISSN 10972765. doi: 10.1016/j.molcel.2022.12.032.

23. Arnaud R. Krebs. Studying transcription factor function in the genome at molecular resolution. Trends in genetics: TIG, 37(9):798–806, September 2021. ISSN 0168-9525. doi: 10.1016/j.tig.2021.03.008.

24. Liesbeth Minnoye, Georgi K. Marinov, Thomas Krausgruber, Lixia Pan, Alexandre P. Marand, Stefano Secchia, William J. Greenleaf, Eileen E. M. Furlong, Keji Zhao, Robert J. Schmitz, Christoph Bock, and Stein Aerts. Chromatin accessibility profiling methods. Nature Reviews. Methods Primers, 1:10, 2021. ISSN 2662-8449. doi: 10.1038/s43586-020-00008-9.

25. Sergio Martin Espinola, Markus Götz, Maelle Bellec, Olivier Messina, Jean-Bernard Fiche, Christophe Houbron, Matthieu Dejean, Ingolf Reim, Andrés M. Cardozo Gizzi, Mounia Lagha, and Marcelo Nollmann. Cis-regulatory chromatin loops arise before TADs and gene activation, and are independent of cell fate during early Drosophila development. Nature Genetics, 53(4):477–486, April 2021. ISSN 1546-1718. doi: 10.1038/s41588-021-00816-z.

26. Yad Ghavi-Helm, Felix A. Klein, Tibor Pakozdi, Lucia Ciglar, Daan Noordermeer, Wolfgang Huber, and Eileen E. M. Furlong. Enhancer loops appear stable during development and are associated with paused polymerase. Nature, 512(7512):96–100, August 2014. ISSN 1476-4687. doi: 10.1038/nature13417.

27. Wouter de Laat and Denis Duboule. Topology of mammalian developmental enhancers and their regulatory landscapes. Nature, 502(7472):499–506, October 2013. ISSN 1476-4687. doi: 10.1038/nature12753.

28. Tim Pollex, Adam Rabinowitz, Maria Cristina Gambetta, Raquel Marco-Ferreres, Rebecca R. Viales, Aleksander Jankowski, Christoph Schaub, and Eileen E. M. Furlong. Enhancer-promoter interactions become more instructive in the transition from cell-fate specification to tissue differentiation. Nature Genetics, 56(4):686–696, April 2024. ISSN 1546-1718. doi: 10.1038/s41588-024-01678-x.

29. Paul W. Hook and Winston Timp. Beyond assembly: the increasing flexibility of singlemolecule sequencing technology. Nature Reviews. Genetics, 24(9):627–641, September 2023. ISSN 1471-0064. doi: 10.1038/s41576-023-00600-1.

30. Can Sönmezer, Rozemarijn Kleinendorst, Dilek Imanci, Guido Barzaghi, Laura Villacorta, Dirk Schübeler, Vladimir Benes, Nacho Molina, and Arnaud Regis Krebs. Molecular Cooccupancy Identifies Transcription Factor Binding Cooperativity In Vivo. Molecular Cell, 81 (2):255–267.e6, January 2021. ISSN 10972765. doi: 10.1016/j.molcel.2020.11.015.

31. Elisa Kreibich, Rozemarijn Kleinendorst, Guido Barzaghi, Sarah Kaspar, and Arnaud R. Krebs. Single-molecule footprinting identifies context-dependent regulation of enhancers by DNA methylation. Molecular Cell, 83(5):787–802.e9, March 2023. ISSN 10972765. doi: 10.1016/j.molcel.2023.01.017.

32. Nour J Abdulhay, Colin P McNally, Laura J Hsieh, Sivakanthan Kasinathan, Aidan Keith, Laurel S Estes, Mehran Karimzadeh, Jason G Underwood, Hani Goodarzi, Geeta J Narlikar, and Vijay Ramani. Massively multiplex single-molecule oligonucleosome footprinting. eLife, 9:e59404, December 2020. ISSN 2050-084X. doi: 10.7554/eLife.59404.

33. Sofia Battaglia, Kevin Dong, Jingyi Wu, Zeyu Chen, Fadi J. Najm, Yuanyuan Zhang, Molly M. Moore, Vivian Hecht, Noam Shoresh, and Bradley E. Bernstein. Long-range phasing of dynamic, tissue-specific and allele-specific regulatory elements. Nature Genetics, 54(10): 1504–1513, October 2022. ISSN 1546-1718. doi: 10.1038/s41588-022-01188-8.

34. Isac Lee, Roham Razaghi, Timothy Gilpatrick, Michael Molnar, Ariel Gershman, Norah Sadowski, Fritz J. Sedlazeck, Kasper D. Hansen, Jared T. Simpson, and Winston Timp. Simultaneous profiling of chromatin accessibility and methylation on human cell lines with nanopore sequencing. Nature Methods, 17(12):1191–1199, December 2020. ISSN 1548-7105. doi: 10.1038/s41592-020-01000-7.

35. Zohar Shipony, Georgi K. Marinov, Matthew P. Swaffer, Nicholas A. Sinnott-Armstrong, Jan M. Skotheim, Anshul Kundaje, and William J. Greenleaf. Long-range single-molecule mapping of chromatin accessibility in eukaryotes. Nature Methods, 17(3):319–327, March 2020. ISSN 1548-7105. doi: 10.1038/s41592-019-0730-2.

36. Andrew B. Stergachis, Brian M. Debo, Eric Haugen, L. Stirling Churchman, and John A. Stamatoyannopoulos. Single-molecule regulatory architectures captured by chromatin fiber sequencing. Science (New York, N.Y.), 368(6498):1449–1454, June 2020. ISSN 1095-9203. doi: 10.1126/science.aaz1646.

37. Rozemarijn W. D. Kleinendorst, Guido Barzaghi, Mike L. Smith, Judith B. Zaugg, and Arnaud R. Krebs. Genome-wide quantification of transcription factor binding at single-DNA-molecule resolution using methyl-transferase footprinting. Nature Protocols, 16(12):5673– 5706, December 2021. ISSN 1750-2799. doi: 10.1038/s41596-021-00630-1.

38. Mark Boltengagen, Daan Verhagen, Michael Roland Wolff, Elisa Oberbeckmann, Matthias Hanke, Ulrich Gerland, Philipp Korber, and Felix Mueller-Planitz. A single fiber view of the nucleosome organization in eukaryotic chromatin. Nucleic Acids Research, 52(1):166–185, January 2024. ISSN 1362-4962. doi: 10.1093/nar/gkad1098.

39. Mitchell R. Vollger, Elliott G. Swanson, Shane J. Neph, Jane Ranchalis, Katherine M. Munson, Ching-Huang Ho, Y. H. Hank Cheng, Adriana E. Sedeño-Cortés, William E. Fondrie, Stephanie C. Bohaczuk, Maxwell A. Dippel, Yizi Mao, Nancy L. Parmalee, Benjamin J. Mallory, William T. Harvey, Younjun Kwon, Gage H. Garcia, Kendra Hoekzema, Jeffrey G. Meyer, Mine Cicek, Evan E. Eichler, William S. Noble, Daniela M. Witten, James T. Bennett, John P. Ray, and Andrew B. Stergachis. A haplotype-resolved view of human gene regulation. bioRxiv, 2025. doi: 10.1101/2024.06.14.599122. Publisher: Cold Spring Harbor Laboratory _eprint: https://www.biorxiv.org/content/early/2025/06/02/2024.06.14.599122.full.pdf.

40. Stephen Small and David N. Arnosti. Transcriptional Enhancers in Drosophila. Genetics, 216(1):1–26, September 2020. ISSN 1943-2631. doi: 10.1534/genetics.120.301370.

41. Arnaud R. Krebs, Dilek Imanci, Leslie Hoerner, Dimos Gaidatzis, Lukas Burger, and Dirk Schübeler. Genome-wide Single-Molecule Footprinting Reveals High RNA Polymerase II Turnover at Paused Promoters. Molecular Cell, 67(3):411–422.e4, August 2017. ISSN 10972765. doi: 10.1016/j.molcel.2017.06.027.

42. Cosmas D. Arnold, Daniel Gerlach, Christoph Stelzer, Łukasz M. Boryń, Martina Rath, and Alexander Stark. Genome-wide quantitative enhancer activity maps identified by STARRseq. Science (New York, N.Y.), 339(6123):1074–1077, March 2013. ISSN 1095-9203. doi: 10.1126/science.1232542.

43. Valentina Baderna, Guido Barzaghi, Rozemarijn Kleinendorst, Kasit Chatsirisupachai, Laura Moniot-Perron, Meike Schopp, Tino Hochepied, Claude Libert, Duncan T Odom, Judith B Zaugg, and Arnaud Krebs. Cumulative TF binding and H3K27 Acetylation drive enhancer activation frequency. bioRxiv, 2025. doi: 10.1101/2025.03.26.645413. Publisher: Cold Spring Harbor Laboratory _eprint: https://www.biorxiv.org/content/early/2025/03/26/2025.03.26.645413.full.pdf.

44. Xiao Li, Xiaona Tang, Xinyang Bing, Christopher Catalano, Taibo Li, Gabriel Dolsten, Carl Wu, and Michael Levine. GAGA-associated factor fosters loop formation in the Drosophila genome. Molecular Cell, 83(9):1519–1526.e4, May 2023. ISSN 1097-4164. doi: 10.1016/j.molcel.2023.03.011.

45. J. Lesley Brown, Liangliang Zhang, Pedro P. Rocha, Judith A. Kassis, and Ming-An Sun. Polycomb protein binding and looping in the ON transcriptional state. Science Advances, 10(17):eadn1837, April 2024. ISSN 2375-2548. doi: 10.1126/sciadv.adn1837.

46. A. Marieke Oudelaar, James O. J. Davies, Lars L. P. Hanssen, Jelena M. Telenius, Ron Schwessinger, Yu Liu, Jill M. Brown, Damien J. Downes, Andrea M. Chiariello, Simona Bianco, Mario Nicodemi, Veronica J. Buckle, Job Dekker, Douglas R. Higgs, and Jim R. Hughes. Single-allele chromatin interactions identify regulatory hubs in dynamic compartmentalized domains. Nature Genetics, 50(12):1744–1751, December 2018. ISSN 1546-1718. doi: 10.1038/s41588-018-0253-2.

47. Mario Iurlaro, Michael B. Stadler, Francesca Masoni, Zainab Jagani, Giorgio G. Galli, and Dirk Schübeler. Mammalian SWI/SNF continuously restores local accessibility to chromatin. Nature Genetics, 53(3):279–287, March 2021. ISSN 1546-1718. doi: 10.1038/s41588-020-00768-w.

48. Nicholas C. Lammers, Yang Joon Kim, Jiaxi Zhao, and Hernan G. Garcia. A matter of time: Using dynamics and theory to uncover mechanisms of transcriptional bursting. Current Opinion in Cell Biology, 67:147–157, December 2020. ISSN 09550674. doi: 10.1016/j.ceb.2020.08.001.

49. Joseph V. W. Meeussen and Tineke L. Lenstra. Time will tell: comparing timescales to gain insight into transcriptional bursting. Trends in genetics: TIG, 40(2):160–174, February 2024. ISSN 0168-9525. doi: 10.1016/j.tig.2023.11.003.

50. Won-Ki Cho, Jan-Hendrik Spille, Micca Hecht, Choongman Lee, Charles Li, Valentin Grube, and Ibrahim I. Cisse. Mediator and RNA polymerase II clusters associate in transcription-dependent condensates. Science (New York, N.Y.), 361(6400):412–415, July 2018. ISSN 1095-9203. doi: 10.1126/science.aar4199.

51. Shasha Chong, Claire Dugast-Darzacq, Zhe Liu, Peng Dong, Gina M. Dailey, Claudia Cattoglio, Alec Heckert, Sambashiva Banala, Luke Lavis, Xavier Darzacq, and Robert Tjian. Imaging dynamic and selective low-complexity domain interactions that control gene transcription. Science (New York, N.Y.), 361(6400):eaar2555, July 2018. ISSN 1095-9203. doi: 10.1126/science.aar2555.

52. Patrick Cramer. Organization and regulation of gene transcription. Nature, 573(7772):45– 54, September 2019. ISSN 1476-4687. doi: 10.1038/s41586-019-1517-4.

53. Bomyi Lim and Michael S. Levine. Enhancer-promoter communication: hubs or loops? Current Opinion in Genetics & Development, 67:5–9, April 2021. ISSN 1879-0380. doi: 10.1016/j.gde.2020.10.001.

54. Benjamin R. Sabari, Alessandra Dall’Agnese, Ann Boija, Isaac A. Klein, Eliot L. Coffey, Krishna Shrinivas, Brian J. Abraham, Nancy M. Hannett, Alicia V. Zamudio, John C. Manteiga, Charles H. Li, Yang E. Guo, Daniel S. Day, Jurian Schuijers, Eliza Vasile, Sohail Malik, Denes Hnisz, Tong Ihn Lee, Ibrahim I. Cisse, Robert G. Roeder, Phillip A. Sharp, Arup K. Chakraborty, and Richard A. Young. Coactivator condensation at super-enhancers links phase separation and gene control. Science (New York, N.Y.), 361(6400):eaar3958, July 2018. ISSN 1095-9203. doi: 10.1126/science.aar3958.

55. Jin H. Yang and Anders S. Hansen. Enhancer selectivity in space and time: from enhancer-promoter interactions to promoter activation. Nature Reviews. Molecular Cell Biology, 25 (7):574–591, July 2024. ISSN 1471-0080. doi: 10.1038/s41580-024-00710-6.

56. Philippe J. Batut, Xin Yang Bing, Zachary Sisco, João Raimundo, Michal Levo, and Michael S. Levine. Genome organization controls transcriptional dynamics during development. Science (New York, N.Y.), 375(6580):566–570, February 2022. ISSN 1095-9203. doi: 10.1126/science.abi7178.

57. Shyam Ramasamy, Abrar Aljahani, Magdalena A. Karpinska, T. B. Ngoc Cao, Taras Velychko, J. Neos Cruz, Michael Lidschreiber, and A. Marieke Oudelaar. The Mediator complex regulates enhancer-promoter interactions. Nature Structural & Molecular Biology, 30(7): 991–1000, July 2023. ISSN 1545-9985. doi: 10.1038/s41594-023-01027-2.

58. Laila El Khattabi, Haiyan Zhao, Jens Kalchschmidt, Natalie Young, Seolkyoung Jung, Peter Van Blerkom, Philippe Kieffer-Kwon, Kyong-Rim Kieffer-Kwon, Solji Park, Xiang Wang, Jordan Krebs, Subhash Tripathi, Noboru Sakabe, Débora R. Sobreira, Su-Chen Huang, Suhas S. P. Rao, Nathanael Pruett, Daniel Chauss, Erica Sadler, Andrea Lopez, Marcelo A. Nóbrega, Erez Lieberman Aiden, Francisco J. Asturias, and Rafael Casellas. A Pliable Mediator Acts as a Functional Rather Than an Architectural Bridge between Promoters and Enhancers. Cell, 178(5):1145–1158.e20, August 2019. ISSN 1097-4172. doi: 10.1016/j.cell.2019.07.011.

59. Karine Lapouge, Rozemarijn Kleinendorst, Kasit Chatsirisupachai, Nikolay Dobrev, Kim Remans, and Arnaud Krebs. Expression, purification and characterization of the GpC methyltransferase M.CviPI v1. February 2023. doi: 10.17504/protocols.io.eq2ly776plx9/v1. https://www.protocols.io/view/expression-purification-and-characterization-of-th-cirbud2n.

60. Jaime A. Castro-Mondragon, Rafael Riudavets-Puig, Ieva Rauluseviciute, Roza Berhanu Lemma, Laura Turchi, Romain Blanc-Mathieu, Jeremy Lucas, Paul Boddie, Aziz Khan, Nicolás Manosalva Pérez, Oriol Fornes, Tiffany Y. Leung, Alejandro Aguirre, Fayrouz Hammal, Daniel Schmelter, Damir Baranasic, Benoit Ballester, Albin Sandelin, Boris Lenhard, Klaas Vandepoele, Wyeth W. Wasserman, François Parcy, and Anthony Mathelier. JASPAR 2022: the 9th release of the open-access database of transcription factor binding profiles. Nucleic Acids Research, 50(D1):D165–D173, January 2022. ISSN 1362-4962. doi: 10.1093/nar/gkab1113.

61. Matthew T. Weirauch, Ally Yang, Mihai Albu, Atina G. Cote, Alejandro Montenegro-Montero, Philipp Drewe, Hamed S. Najafabadi, Samuel A. Lambert, Ishminder Mann, Kate Cook, Hong Zheng, Alejandra Goity, Harm van Bakel, Jean-Claude Lozano, Mary Galli, Mathew G. Lewsey, Eryong Huang, Tuhin Mukherjee, Xiaoting Chen, John S. Reece-Hoyes, Sridhar Govindarajan, Gad Shaulsky, Albertha J. M. Walhout, François-Yves Bouget, Gunnar Ratsch, Luis F. Larrondo, Joseph R. Ecker, and Timothy R. Hughes. Determination and inference of eukaryotic transcription factor sequence specificity. Cell, 158(6):1431–1443, September 2014. ISSN 1097-4172. doi: 10.1016/j.cell.2014.08.009.

62. Hervé Pagès, Patrick Aboyoun, Robert Gentleman, and Saikat DebRoy. Biostrings: Efficient manipulation of biological strings, 2024.

63. Raivo Kolde. pheatmap: Pretty Heatmaps, 2019.

64. F. Krueger, F. James, P. Ewels, E. Afyounian, and B. Schuster-Boeckler. FelixKrueger/TrimGalore, v0.6.7 via Zenodo., 2021.

65. Dimos Gaidatzis, Anita Lerch, Florian Hahne, and Michael B. Stadler. QuasR: quantification and annotation of short reads in R. Bioinformatics, 31(7):1130–1132, April 2015. ISSN 1367-4811, 1367-4803. doi: 10.1093/bioinformatics/btu781.

66. Ben Langmead, Cole Trapnell, Mihai Pop, and Steven L Salzberg. Ultrafast and memory-efficient alignment of short DNA sequences to the human genome. Genome Biology, 10(3): R25, 2009. ISSN 1465-6906. doi: 10.1186/gb-2009-10-3-r25.

67. Broad Institute. Picard Toolkit, Broad Institute, GitHub Repository, 2019.

68. Alexander Dobin, Carrie A. Davis, Felix Schlesinger, Jorg Drenkow, Chris Zaleski, Sonali Jha, Philippe Batut, Mark Chaisson, and Thomas R. Gingeras. STAR: ultrafast universal RNA-seq aligner. Bioinformatics (Oxford, England), 29(1):15–21, January 2013. ISSN 1367-4811. doi: 10.1093/bioinformatics/bts635.

69. Yang Liao, Gordon K. Smyth, and Wei Shi. The R package Rsubread is easier, faster, cheaper and better for alignment and quantification of RNA sequencing reads. Nucleic Acids Research, 47(8):e47, May 2019. ISSN 1362-4962. doi: 10.1093/nar/gkz114.

70. Arzu Öztürk Çolak, Steven J. Marygold, Giulia Antonazzo, Helen Attrill, Damien Goutte-Gattat, Victoria K. Jenkins, Beverley B. Matthews, Gillian Millburn, Gilberto Dos Santos, Christopher J. Tabone, and FlyBase Consortium. FlyBase: updates to the Drosophila genes and genomes database. Genetics, 227(1):iyad211, May 2024. ISSN 1943-2631. doi: 10.1093/genetics/iyad211.

71. Michael I. Love, Wolfgang Huber, and Simon Anders. Moderated estimation of fold change and dispersion for RNA-seq data with DESeq2. Genome Biology, 15(12):550, 2014. ISSN 1474-760X. doi: 10.1186/s13059-014-0550-8.

